# High fat diet worsens pathology and impairment in an Alzheimer’s mouse model, but not by synergistically decreasing cerebral blood flow

**DOI:** 10.1101/2019.12.16.878397

**Authors:** Oliver Bracko, Lindsay K. Vinarcsik, Jean C. Cruz Hernández, Nancy E. Ruiz-Uribe, Mohammad Haft-Javaherian, Kaja Falkenhain, Egle M. Ramanauskaite, Muhammad Ali, Aditi Mohapatra, Madisen Swallow, Brendah N. Njiru, Victorine Muse, Stall Catchers contributors, Pietro E. Michelucci, Nozomi Nishimura, Chris B. Schaffer

**Affiliations:** Meinig School of Biomedical Engineering, Cornell University, Ithaca, NY, USA; Human Computation Institute, Ithaca, NY, USA

**Author notes:** Worldwide, list of individual Stall Catchers contributors who contributed to this study is at https://hcinst.org/sc-hfd.

**Keywords:** animal behavior, nonlinear microscopy, brain blood flow, western high fat diet

## Abstract

Obesity is linked to increased risk for and severity of Alzheimer’s disease (AD). Cerebral blood flow (CBF) reductions are an early feature of AD and are also linked to obesity. We showed that non-flowing capillaries, caused by adhered neutrophils, underlie the CBF reduction in mouse models of AD. Because obesity could exacerbate the vascular inflammation likely underlying this neutrophil adhesion, we tested links between obesity and AD by feeding APP/PS1 mice a high fat diet (Hfd) and evaluating behavioral, physiological, and pathological changes. We found trends toward poorer memory performance in APP/PS1 mice fed a Hfd, impaired social interactions with either APP/PS1 genotype or a Hfd, and synergistic impairment of sensory-motor function in APP/PS1 mice fed a Hfd. The Hfd led to increases in amyloid-beta monomers and plaques in APP/PS1 mice, as well as increased brain inflammation. These results agree with previous reports showing obesity exacerbates AD-related pathology and symptoms in mice. We used a crowd-sourced, citizen science approach to analyze imaging data to determine the impact of the APP/PS1 genotype and a Hfd capillary stalling and CBF. Surprisingly, we did not see an increase in the number of non-flowing capillaries or a worsening of the CBF deficit in APP/PS1 mice fed a Hfd as compared to controls, suggesting capillary stalling is not a mechanistic link between a Hfd and increased severity of AD in mice. Reducing capillary stalling by blocking neutrophil adhesion improved CBF and short-term memory function in APP/PS1 mice, even when fed a Hfd.

**Significance statement:** Obesity, especially in mid-life, has been linked to increased risk for and severity of Alzheimer’s disease. Here, we show that blocking adhesion of white blood cells leads to increases in brain blood flow that improve cognitive function, regardless of whether mice are obese or not.

## Introduction

Recent evidence has suggested that the presence of cardiovascular and metabolic risk factors [1, 2] during midlife increases the risk for and severity of late-life dementia. For example, midlife diabetes [3], smoking [4], hypertension [4], and hyperlipidemia [5] all increase the risk for late-life dementia in a dose-dependent manner [5–7]. In addition to being associated with increased general morbidity and mortality, midlife obesity is also a strong predictor of dementia incidence and severity [8, 9]. Thus, the worldwide increasing rates of overweightness and obesity present a major challenge for chronic disease prevention, including neurodegenerative diseases such as Alzheimer’s disease (AD) [10, 11].

Feeding mouse models of neurodegenerative disease with diets that induce obesity is one approach to elucidating the mechanistic links between obesity and dementia. Feeding mice a western high-fat diet (Hfd) has been linked to impaired cognitive function in wild-type mice [12, 13], as well as to more severe cognitive impairment in AD mouse models [14, 15], with obesity-linked cognitive deficits most prominent in tests for short-term and working memory [16, 17]. For example, a recent study by Walker, et al. showed more severe deficits in the Y-maze and nest building behavioral assays in the APP/PS1 mouse model of AD when fed a Hfd for several months, as compared to APP/PS1 mice eating normal chow [18]. The severity of the impacts of a Hfd on cognitive function in neurodegenerative disease models that have been reported vary widely, likely due to different diets, mouse models, starting ages, and durations of the Hfd treatment [19–21]. In a recent report by Salas, et al., the authors provide a summary of different diet parameters and AD mouse models that have been explored and the different cognitive outcomes observed [19].

Early studies to identify potential mechanisms underlying the exacerbation of cognitive dysfunction in AD mouse models fed a Hfd focused on alterations in the pathological hallmarks of AD. The emerging picture from this body of work is that a Hfd leads to increased deposition of amyloid-beta (Aβ) as plaques in the brain, as well as to increased brain inflammation, likely driving more Aβ-and inflammation-induced neural dysfunction. However, there have been variable findings on the impact of Hfd on the number and composition of amyloid-beta (Aβ) aggregates, the amount of hyperphosphorylated tau, and the degree of brain inflammation [18, 22, 23], again likely due to differences in diet parameters and AD mouse models. More recently, other mechanisms that may link obesity and consumption of a Hfd to dementia have been explored. Synaptic pruning by activated microglia was found to be elevated in obese rodents [12, 24, 25], and such rewiring of neural connections could contribute to Hfd-associated cognitive impairment. Recent work has also shown that a Hfd induced changes in the organismal diversity of the gut microbiome that was specifically linked to changes in inflammatory signaling within the hippocampus as well as to deficits in short-term memory [26].

Obesity has been linked to decreases in CBF [27–29], which could be implicated in the increased incidence and severity of AD in the obese population [30, 31]. We have recently shown that circulating neutrophils transiently adhere in a small fraction of cortical capillary segments in APP/PS1 mice. These capillary stalls were released by administering an antibody against the neutrophil specific protein Ly6G, leading to a ~20% increase in cerebral blood flow (CBF), suggesting this capillary plugging underlies the CBF deficit in this AD mouse model. This CBF increase led to improved performance on spatial and working memory tasks in APP/PS1 mice, even at late stages of disease progression [32, 33]. Obesity is also a driver of chronic, systemic inflammation. For example, studies in both humans [34] and mouse models [35, 36] have shown that obesity is linked to increased secretion of plasma fatty acids from adipose tissue into the vascular system, which causes inflammatory responses in the periphery as well as in the central nervous system[37]. This systemic inflammation could drive activity by circulating immune and inflammatory cells that may contribute to neurological dysfunction. For example, Hfd-fed mice show increased invasion of monocytes into the central nervous system compared to animals on a normal diet [38]. We hypothesized that a Hfd might increase the severity of vascular inflammation in the brain and exacerbate the capillary stalling and CBF deficits seen in APP/PS1 mice. To test this idea, we fed APP/PS1 and WT mice a Hfd or normal chow and assessed the impact on cognitive and sensory-motor function as well as Aβ deposition and brain inflammation. We then used *in vivo* two-photon excited fluorescence (2PEF) microscopy to quantify cortical blood flow and the incidence of capillary stalling in these animals.

## Methods

### Animals, diet, and experimental groups

All animal procedures were approved by the Cornell Institutional Animal Care and Use Committee and were performed under the guidance of the Cornell Center for Animal Resources and Education. We used adult transgenic mice (APP/PS1; B6.Cg-Tg (APPswe, PSEN1dE9) 85Dbo/J; RRID: MMRRC_034832-JAX), The Jackson Laboratory) as a mouse model of AD and littermate wild-type (WT) mice (C57BL/6) as controls. Starting at three months of age, half of the APP/PS1 and WT mice were switched from standard lab chow (Teklad LM-485, Envigo) to a high-fat diet (Hfd, TD.88137 from Harlan). The Hfd had 42% of food calories derived from fat, 42.7% from carbohydrates, and 15.2% from protein sources, while the standard chow had 13% from fat, 58% from carbohydrates, and 29% from protein. The mice were weighed weekly throughout the entire experiment until they were sacrificed for histological analysis. We compared four groups with the following number of mice in each group — wild-type (WT) mice on standard chow (WT-NC): n=10; WT mice on Hfd (WT-Hfd): n=15; APP/PS1 mice on standard chow (APP/PS1-NC): n=10; and APP/PS1 mice on Hfd (APP/PS1-Hfd): n=15. All animals underwent behavioral testing at ~8 and ~10 months of age. After testing at 10 months of age, three animals from each group received cranial windows and were imaged to examine capillary stalling and cerebral blood flow after 3-4 weeks of surgical recovery. The remaining animals underwent additional behavioral testing at 15 and 19 months of age. After the behavioral testing at 19 months, mice received a single treatment with an antibody against Ly6G (α-Ly6G; IP 4 mg/kg; BD Bioscience No.: 561005) or with an isotype control antibody (Iso-Ctr; BD Bioscience No.: 553929) and behavioral testing was repeated one day after antibody administration. All animals then received cranial windows and were imaged, again after 3-4 weeks of recovery, to study cortical hemodynamics before and after a second treatment with α-Ly6G or Iso-Ctr antibodies at ~21 months of age. After the last imaging session, animals were perfused for molecular and histopathological analysis of brain tissue. See experimental timeline in Fig. 1A. Several animals were excluded from the 21 month imaging studies because the cranial window was not clear enough for high quality optical imaging. In our study, there was a greater tendency for animals on the Hfd to develop these cloudy windows (Number of animals that were excluded from 21 month imaging — APP/PS1-NC: n=2; WT-Hfd: n=4, and APP/PS1-Hfd= n=3). Whether or not the animal could be imaged, the brain was still collected for molecular and histopathological analysis.

**Figure 1:**
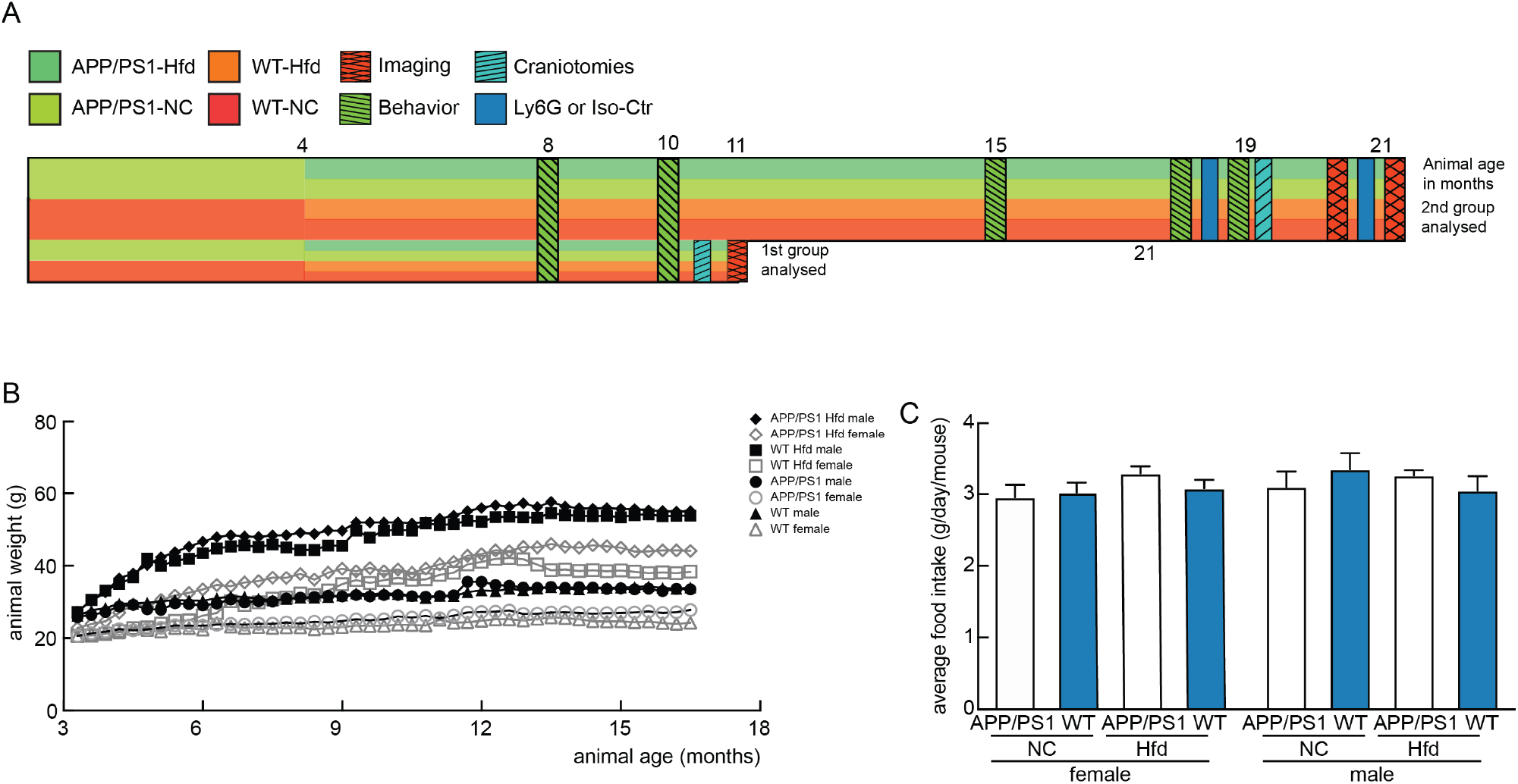
Experimental design/timeline, and animal weight gain and food consumption. (A) Schematic illustrating animal groups, timeline, measurements, and interventions. APP/PS1 and WT mice were allocated to a high fat diet (Hfd) or normal chow (NC) at 4 months of age. Animals received behavioral testing multiple times, as indicated. A small cohort from all four groups received cranial windows and cortical imaging at 11 months of age. The remaining cohort received additional behavioral testing, including before and after administration of α-Ly6G at 19 months, and then received cranial windows and were imaged before and after α-Ly6G treatment. All experimental groups were kept on their diet continuously from four months of age until euthanasia. (B) Average mouse weight for each group as a function of animal age, beginning at 4 months of age, distinguishing male and female mice in each group. (C) Average mass of food intake per mouse per day for all groups, distinguishing female (left) and male (right) animals.

### Behavior Experiments

Animals were taken into the behavior room one hour prior to the experiment to allow accommodation. All experiments were performed under red light in an isolated room and the position of the mouse’s nose was automatically traced by Viewer III software (Biobserve, Bonn, Germany). In addition to the automatic results obtained by Viewer III software, a blinded experimenter independently scored all mouse behavior manually. Behavioral analysis of untreated animals was conducted at multiple time points and, at 19 months of age, at one day after IP administration of α-Ly6G or Iso-Ctr antibodies. The experimenter was blinded with regard to the genotype and diet of the mice, although it was easy to distinguish the Hfd-fed mice at many timepoints due to their weight gain.

### Object replacement test

The object replacement (OR) task evaluated spatial memory performance. All objects were validated in a separate cohort of mice to ensure that no intrinsic preference or aversion was observed and animals explored all objects similarly. Exploration time for the objects was defined as any time when there was physical contact with an object (whisking, sniffing, rearing on or touching the object) or when the animal was oriented toward the object and the head was within 2 cm of the object. In trial 1, mice were allowed to explore two identical objects for 10 min in the arena and then returned to their home cage for 60 min. Mice were then returned to the testing arena for 3 min with one object moved to a novel location (trial 2). Care was taken to ensure that the change of placement altered both the intrinsic relationship between objects (e.g. a rotation of the moved object) and the position relative to internal visual cues (one wall of testing arena had a pattern) (e.g. new location in the arena). The preference score (%) for OR tasks was calculated as ([exploration time of the novel object]/[exploration time of both objects]) × 100 from the data in trial 2. Automated tracking and manual scoring yielded similar results across groups, so we report the automated tracking results.

### Y-Maze

The Y-Maze task was used to measure working memory by quantifying spontaneous alternation between arms of the maze. The Y-maze consisted of three arms at 120° and was made of light grey plastic. Each arm was 6-cm wide and 36-cm long, and had 12.5-cm high walls. The maze was cleaned with 70% ethanol after each mouse. A mouse was placed in the Y-maze and allowed to explore for 6 min. Mouse behavior was monitored, recorded, and analyzed using the Viewer software. A mouse was considered to have entered an arm if the whole body (except for the tail) entered the arm and to have exited if the whole body (except for the tail) exited the arm. If an animal consecutively entered three different arms, it was counted as an alternating trial. Because the maximum number of triads (three successive alternating arm entries) is the total number of arm entries minus 2, the spontaneous alternation score was calculated as (the number of alternating triads)/(the total number of arm entries - 2) × 100.

### Balance beam walk

Motor coordination and balance of mice were assessed by measuring the ability of the mice to traverse a graded series of narrow beams to reach an enclosed safety platform. The beams consisted of 80-cm long strips of round wood with 12-and 6-mm diameters. The beams were placed horizontally, 40 cm above the floor, with one end mounted on a narrow support and the other end attached to an enclosed platform in which the mice could hide. A bright light was shined on the narrow support at the beginning of the beam. Mice received three consecutive trials on each of the round beams, in each case progressing from the widest to the narrowest beam (15 min between each trail). Mice were allowed up to 60 s to traverse each beam. The time required to traverse each beam and the number of times the hind feet slipped off each beam were recorded for each trial. Analysis of each measure was based on the mean of the last two trials for each beam (mice typically take one trial to habituate to the task, so the first trial for each beam was excluded). There were no notable deficits with the 12-mm diameter beam, so we report results only for the 6-mm diameter beam.

### Social interaction test

The test for sociability and preference for social novelty was conducted as previously described [39]. The apparatus comprised a rectangular, three-chambered box. Each chamber was 20 × 20 × 22 cm and the dividing walls were made from clear Plexiglas, with small square openings (6 × 4 cm) allowing access into each chamber. An unfamiliar C57BL/6J mouse of the same sex and age as the subject mouse (stranger 1) that had no prior contact with the subject mouse was placed in one of the side chambers. The placement of stranger 1 in the left or right-side chambers was systematically alternated between trials. The stranger mouse was enclosed in a small, 8 × 5 × 4 cm wire cage that allowed nose contact between the bars but prevented fighting. The subject mouse was first placed in the middle chamber and allowed to explore the arena for 5 min. Then the door was opened, and the mouse could freely explore all chambers including the one with the stranger for 10 min. More sociable mice will spend more time with the stranger, which was quantified in two ways. First, we defined the direct contact time as the fraction of time the subject mouse spends within 3 cm of the stranger and with their nose oriented toward the stranger. Second, we defined the chamber time as the fraction of time the subject mouse spends in the chamber that contains the stranger. At the end of the first 10 min, each subject mouse was tested in a second 10 min session to quantify social preference for a new stranger. A second, unfamiliar mouse of the same age and sex as the subject mouse (stranger 2) was placed in the chamber that had been empty during the first 10 min session, again enclosed in a small wire box. The subject mouse then had a choice between the first, already-investigated unfamiliar mouse (stranger 1), and the novel unfamiliar mouse (stranger 2). The direct contact time and chamber time were quantified, as above, for both strangers. The stranger mice used in this experiment were age matched C57BL/6J mice that were not littermates to the subject mice. All stranger mice were on standard chow. Stranger mice were alternated between trials, with each stranger participating in 3-5 social interaction assays for different subject mice.

### Surgical preparation and in vivo imaging of blood flow and vessel diameter

The surgery and in vivo two-photon microscopy were performed as previously described [32]. In brief, a 6-mm diameter craniotomy was prepared over parietal cortex and covered by gluing a glass coverslip to the skull. Animals rested for at least 3-4 weeks before imaging to minimize the impact of post-surgical inflammation. During the imaging session, mice were placed on a custom stereotactic frame and were anesthetized using ~1.5% isoflurane in 100% oxygen, with small adjustments to the isoflurane level made to maintain the respiratory rate at ~1 Hz. Mice were kept at 37 °C with a feedback-controlled heating pad (40-90-8D DC, FHC). After being anesthetized and once an hour during the imaging session mice received a sub-cutaneous injection of atropine (0.005 mg/100 g mouse weight; 54925-063-10, Med Pharmex) to prevent lung secretions and of 5% glucose in saline (1 mL/100 g mouse weight) to prevent dehydration. For imaging, leukocytes and blood platelets were labeled with a retro-orbital injection of Rhodamine 6G (0.1ml, 1mg ml-1 in 0.9% saline, Acros Organics, Pure). Leukocytes were distinguished from blood platelets with a retro-orbital injection of Hoechst 33342 (50 μl, 4.8 mg ml-1 in 0.9% saline, Thermo Fisher Scientific). Texas Red dextran (40 μl, 2.5%, molecular weight = 70,000 kDa, Thermo Fisher Scientific) in saline was also injected retro-orbitally to label the vasculature. Mice were imaged on a locally-designed two-photon excited fluorescence (2PEF) microscope, using 830-nm, femtosecond duration laser pulses for excitation (Vision II, Coherent). Emission was detected on three photomultiplier tubes through the following emission filters: 417/60 nm (center wavelength/bandwidth) for Hoechst, 550/49 nm for Rhodamine 6G, and 641/75 nm for Texas Red. We collected three-dimensional image stacks and identified penetrating arterioles based on their connectivity to readily-identifiable surface arterioles and by determining their flow direction using the motion of unlabeled red blood cells in the sea of fluorescently-labeled blood plasma. In each mouse, we identified 5-15 penetrating arterioles and took small three-dimensional image stacks to determine the diameter as well as line-scans along the centerline of the vessel to determine red blood cell flow speed [40]. Volumetric blood flow, *F*, was calculated as

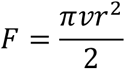

where *v* is the centerline RBC speed and *r* is the vessel radius. These measures were taken at 11 months of age in a set of three mice per group and in the remaining animals at 21 months of age. At the 21 months time point, imaging was conducted at baseline and ~1 day after injection of α-Ly6G or isotype control antibodies (IP 4 mg/kg).

### Post-mortem tissue analysis

After the last imaging session, all animals were sacrificed by lethal injection of pentobarbital (5 mg/100 g). Brains were quickly extracted and divided along the centerline. One half was immersed in 4% paraformaldehyde in phosphate buffered saline (PBS) for later immunohistological analysis and the other half was snap frozen in liquid nitrogen for ELISA, qPCR, and cytokine array assays. The frozen APP/PS1 mouse hemi-brains were weighed and sectioned into 1-mm thick coronal slices, with alternating sections divided across two tubes. This post-mortem tissue analysis was conducted on the following mice — 11 months of age: WT-NC: n=3; WT-Hfd: n=3; APP/PS1-NC: n=3; APP/PS1-Hfd: n=3; 21 months of age: WT-NC: n=4; WT-Hfd: n=4; APP/PS1-NC: n=4; APP/PS1-Hfd: n=4.

### ELISA assay

ELISA assays were used to quantify Aβ1-40 and Aβ1-42 load in soluble and insoluble fractions. Briefly, one series of frozen brain slices was homogenized in 1 ml PBS containing complete protease inhibitor (Roche Applied Science) and 1 mM AEBSF (Sigma) using a Dounce homogenizer. The homogenates were then sonicated and centrifuged at 14,000 g for 30 min at 4° C. The supernatant (soluble fraction) was removed and stored at −80° C. The pellet was redissolved in 0.5 ml 70% formic acid, sonicated, and centrifuged at 14,000 g for 30 min at 4° C, and the supernatant was removed and neutralized using 1M Tris buffer at pH 9 (insoluble fraction). Protein concentration was measured in the soluble fraction and the insoluble fraction using the Pierce BCA Protein Assay (Thermo Fischer Scientific). The soluble fraction extracts were diluted 1:5, while the formic acid extracts were diluted 1:1 after neutralization. These brain extracts were analyzed by sandwich ELISA for Aβ1-40 and Aβ1-42 using commercial ELISA kits and following the manufacturer’s protocol (Aβ1-40: KHB348, and Aβ1-42: KHB3441, Thermo Fisher Scientific). The Aβ concentration was calculated by comparing the sample absorbance with the absorbance of known concentrations of synthetic Aβ1-40 and Aβ1-42 standards on the same plate. Data was acquired with a Synergy HT plate reader (Biotek) and analyzed using Gen5 software (BioTek) and Prism (Graphpad).

### Cytokine array

Some of the soluble fraction that was extracted for ELISA was further analyzed using a 62 target mouse cytokine array (ab133995, Abcam) to examine changes in inflammatory signaling. The soluble fraction was thawed and was then centrifuged at 14,000 g for 30 min at 4° C. The supernatant was collected, and total protein concentration was quantified using a BCA protein assay (Pierce BCA Protein Assay kit, 23225, ThermoFisher Scientific). Samples were diluted to 0.8 ug/uL with a 1X cell lysis buffer provided by the manufacturer. The cytokine assay was performed following the instructions from the manufacturer, and the arrays were read with a ChemBio chemiluminescence reader, using a 2 min. exposure time, which we had determined avoids saturation of the positive control dots on the array. The array data was quantified using the protein array analyzer plugin from ImageJ [41]. All samples were normalized to the reference dots on each array, to express the results as a relative density, which was then compared across animal groups.

### Extraction of total RNA, cDNA synthesis, and RT-qPCR

We examined the impact of a Hfd on the activity of the murine APP gene using the second series of snap frozen slices. After cutting, these slices were immediately transferred to 0.5 ml of RNA stabilization solution (RNAlater, ThermoFischer Scientific). Total RNA was isolated from about 50 mg of sample using RNeasy mini Kit (Qiagen) according to the manufacturer’s instructions. After extraction, the RNA concentration was estimated by spectrophotometric analysis using a NanoDrop 1000 (ThermoFischer Scientific). cDNA copies of total RNA were obtained using the High-Capacity RNA-to-cDNA^™^ Kit (ThermoFischer Scientific) and then adjusted to a concentration equivalent to 100 ng/μl of total RNA using nuclease-free water (ThermoFischer Scientific). The qPCR Taqman probes were purchased from ThermoFischer Science (mAPP: AssayID Mm01344172_m1; and GAPDH: Assay ID Mm99999915_g1). RT-qPCR was performed with 10 μl cDNA (corresponding to the cDNA reverse transcribed from approximately 10 ng of RNA) in 20 μl of reaction mix containing 10 μl TaqMan Gene Expression Master Mix (2×). qPCR data were acquired with technical triplicates using these settings: 50°C for 2 min; 95°C for 10 min for first cycle, then 30 s for subsequent cycles; 60°C for 1 min; 40 cycles total on an ABI PRISM 7900 HT Sequence Detection System (Applied Biosystems). The threshold cycle (Ct) was calculated by the instrument software automatically. GAPDH was used as an endogenous normalization control and the fold expression relative to GAPDH was determined by the ΔΔCt method.

### Immunohistochemistry

The density and morphology of glial cells, as well as the density of amyloid plaques, was quantified using immunohistochemistry on the half brains that were fixed in PFA. The half brains were soaked in 30% sucrose in PBS until saturated, then frozen in optimal cutting temperature (OCT) compound (Fisher Healthcare) and cut in 30-μm thick sagittal sections. Every sixth section from each mouse was mounted and incubated overnight at 4° C with 1:250 anti-rabbit Iba1 (019-19741, WAKO) and 1:500 anti-chicken-GFAP (ab4674), Abcam). Samples were then incubated with secondary antibodies for two hours at 1:300 (ThermoFischer Scientific). Finally, we counterstained with 1:100 Methoxy-X04 (1 mg/ml MeO-X04 (5 mg/ml in 10% DMSO, 45% propylene glycol, and 45% saline) 4920, Tocris) for 15 min at room temperature and then washed the slides twice with 80% ethanol for 2 min. The sections were mounted using Richard-Allan Scientific^™^ Mounting Medium (4112APG, ThermoFischer Scientific). Images were taken using confocal microscopy (Zeiss Examiner.D1 AXIO) and cortical and hippocampal regions of interest were defined in each section anatomically. Images of Methoxy-X04 were then manually thresholded, using the same threshold for all sections from a given mouse. Appropriate thresholds varied mouse to mouse and were set to ensure that the smallest Methoxy-X04 labeled objects that morphologically appeared to be an amyloid plaque remained above threshold. The fraction of pixels above threshold was determined across all sections for these regions of interest. In addition, the number of amyloid deposits were manually counted as the number of Methoxy-X04 positive objects. For Iba1 and GFAP images, the number of Iba1 and GFAP positive cells were counted using the automated counting tool in ImageJ (NIH). All image processing was done blinded to the experimental group. All sections were stained and imaged in parallel.

### Quantification of flowing and non-flowing vessel segments

We scored about 96,000 individual capillary segments as flowing or stalled for this study. The fluorescent dye we intravenously injected to label the blood plasma generated a negative contrast image of blood cells, so the motion of the dark patches produced by these cells could be used to measure blood flow. We take 1-μm spaced image stacks at ~1 frame per second, so that each capillary segment is visible for a minimum of ~five frames or 5 s. We define capillaries as stalled if we observe a dark patch in the lumen of the vessel that does not move over the observation time for that capillary segment. For this study, this scoring of capillaries as flowing or stalled used a data analysis pipeline that involved the following steps: 1. Curation of image stacks of brain vasculature; 2. Segmentation of those images using a machine learning approach; 3. Identification of capillary segments; 4. Crowd-sourced scoring of individual capillaries as flowing or stalled, and 5. Internal quality control of crowd-sourced results.

1. First the raw image stacks of the vasculature were masked to remove the larger surface vessels and regions where the signal to noise was poor using ImageJ (NIH). Masked images were then preprocessed to normalize the image intensity distribution and spatial resolution [42] and to remove motion artifacts [43].
2. The vasculature network in each image stack was then segmented into binary images using a deep convolutional neural network, DeepVess, we recently developed [42].
3. Individual capillaries, defined as a segment between two bifurcations, were identified by extracting the vasculature centerlines using sequential dilation and thinning operations [42], in addition to some centerline artifact removal steps [42].
4. The “Stall Catchers” platform was designed to allow citizen scientists to score capillary segments as flowing or stalled (www.stallcatchers.com). For each capillary segment, a sub-stack was cropped from the original image stack that was centered on the segment of interest and the image intensity within the cropped sub-stack was normalized to utilize the entire available intensity range. Then, an outline was drawn around the segment of interest based on a dilation of the centerline. The outline’s characteristics were selected based on user feedback gathered from alpha testing. Citizen scientists then used the Stall Catchers interface to view these image stacks. Players could scan up and down in the stack (and therefore forward and backward in time) and they were asked to judge whether they saw moving dark patches in the outlined capillary segment. In addition to image stacks where we did not know if the vessel was flowing or not (“research vessels”), each player also received image stacks where we had already scored the identified capillary (“calibration vessels”) to maintain a running estimate of each player’s sensitivity. The weight of a player’s opinion in the final averaged answer for a capillary increased with this sensitivity. In addition, as players demonstrated greater skill, they were shown a higher proportion of research vessels as they were deemed to need less training and less frequent monitoring. Because the base rate of stalling in research vessels was less than 1%, the base rate of being shown stalled capillary segments is easily manipulated via the calibration vessels. Thus, as the ratio of research vessels to calibration vessels changed due to player skill level, the ratio of flowing to stalled calibration vessels was adjusted to maintain a constant overall ratio of flowing to stalled when including the research vessels. This ratio was kept constant to reduce potential response biases that might otherwise result. Each capillary segment was scored by multiple players and the results were averaged, weighted by each player’s accuracy at the time they scored that vessel, yielding a “crowd confidence” score for each vessel to be stalled, with crowd confidence closer to 1 implying a high probability of being stalled and a crowd confidence of 0 implying almost certainly flowing. We determined enough player answers for a capillary had been collected when an empirically-determined sensitivity threshold was reached, which was simply the sum of the sensitivities of all the individual players that provided an answer for that capillary. This threshold sensitivity was set so that all vessels that were known to be stalled in a test data set were near the top of the rank-ordered crowd confidence score.
5. Researchers in our laboratory then inspected this crowd-sourced scoring, starting with the vessels that had the highest crowd confidence of being stalled and continuing to vessels with a crowd confidence of 50% (about 2% of the total capillary segments). For vessels with a crowd confidence score greater than 90% (277 vessel segments), we found that 100% of those vessels were actually stalled. At a crowd confidence score between 50% and 60% (475 vessel segments), we found that only one of 158 vessels were actually stalled. Reported rates of capillary stalling for each image stack were obtained by taking the researcher-determined capillary segments with crowd confidence greater than 50% that were, in fact, stalled and dividing by the total number of capillary segments in that image stack.

### Data Visualization and Statistical Analysis

In the figures in the main text, longitudinal behavioral data as well as data on capillary stalling and flow speed for different age animals are shown as the average of each experimental group with the SD indicated. This representation simplifies visualization of trends over animal age, between different genotypes and diets, and in response to α-Ly6G treatment. We further show all animal-by-animal behavioral data as well as measurements of changes in vessel diameter, capillary stalls, and blood flow in the supplementary information as boxplots. We use Tukey boxplots, where the box contains the middle two quartiles of the data, the whiskers extend 1.5 times the difference between the 75th and 25th percentiles, the black line defines the median, and the red line defines the mean, which is calculated excluding any outliers that sit outside the whiskers. Data on amyloid concentrations and expression of inflammatory markers in the main figures and supplementary materials is similarly represented with boxplots. For all statistical analysis, normality was tested with the D’Agostino-Pearson test and when all data for a comparison was normally distributed, the statistical significance of differences between groups was evaluated by one-way analysis of variance (ANOVA) followed by pairwise comparisons using the multiple-comparison corrected Holm-Šídák test. When any data was not normally distributed, a Kruskal-Wallis test with multiple comparisons correction was performed. P-values <0.05 were considered statistically significant. All statistical analyses were performed using Prism 7/8 software (GraphPad Software) or JMP Pro 14 (SAS Institute Inc.). We used a standardized set of significance indicators to distinguish different comparisons: * for comparisons between mouse genotype; # for comparisons between treatment groups; † for comparisons between diets; and ^λ^ for comparisons between animal ages.

## Results

APP/PS1 mice and wild-type (WT) mice were fed a high-fat diet (Hfd) starting at 4 months of age. When cognitive deficits first became evident in the APP/PS1 mice, a subset of mice were taken out and imaged using *in vivo* two-photon excitation microscopy (2PEF) for quantification of capillary stalling and cerebral blood flow (CBF) at 11 months of age (Fig. 1A). The remaining mice received additional behavioral testing and were kept on the Hfd until 19 months of age (Fig. 1A), when they were then treated with antibodies against Ly6G (α-Ly6G) or with isotype control (Iso-Ctr) antibodies. These animals then received cranial windows and were imaged before and after treatment with α-Ly6G or control antibodies at 21 months of age. Animals were then sacrificed and histological analysis was performed on the brain tissue extracted post-mortem. We label the four groups we compared as: WT mice on a normal chow (WT-NC), WT mice on the Hfd (WT-Hfd), APP/PS1 mice on the normal chow (APP/PS1-NC), and APP/PS1 mice on the Hfd (APP/PS1-Hfd).

All female mice displayed a slower curve of weight gain compared to male mice on either a normal chow or Hfd. There was no significant difference in weight gain between WT and APP/PS1 mice for either sex on either diet (Fig. 1B). We found no difference in food uptake between all groups, measured as the average food uptake per cage per week and assuming each mouse ate equally (Fig. 1C).

Using the extracted brain tissue, we determined the impact of the Hfd on amyloid burden in the APP/PS1 mice by measuring the levels of Aβ_(1-40)_ and Aβ_(1-42)_ monomers and the density of amyloid plaques from animals sacrificed at 11 and 21 months of age. The amount of Aβ_(1-40)_ in the PBS-soluble fraction was increased at 11 months of age and remained elevated at 21 months of age in APP/PS1-Hfd mice as compared to APP-NC animals (Fig. 2A). A similar tendency was seen in the insoluble fraction of Aβ_(1-40)_ (Fig. 2B). The levels of Aβ_(1-42)_ were similar between APP/PS1-Hfd and APP/PS1-NC mice in both the PBS-soluble and the insoluble fractions at both ages, but with a trend towards increased PBS-soluble Aβ_(1-42)_ at 21 months of age in the APP/PS1-Hfd group (Fig. 2C and D). Next, we analyzed the overall amyloid plaque load in serial sections from cortical and hippocampal regions using Methoxy-X04 staining (Fig. 2E). We found an increase in Methoxy-XO4 positive pixels in APP/PS1-Hfd mice as compared to APP/PS1-NC mice in the cortex at 21 months, but not at 11 months of age (Fig. 2F). In the hippocampus, amyloid plaque density was increased in APP/PS1-Hfd mice, as compared to APP/PS1-NC animals, at both time points (Fig. 2G). Hippocampal, but not cortical, amyloid deposits increased with age in mice fed either normal chow or the Hfd (Fig. 2G).

**Figure 2:**
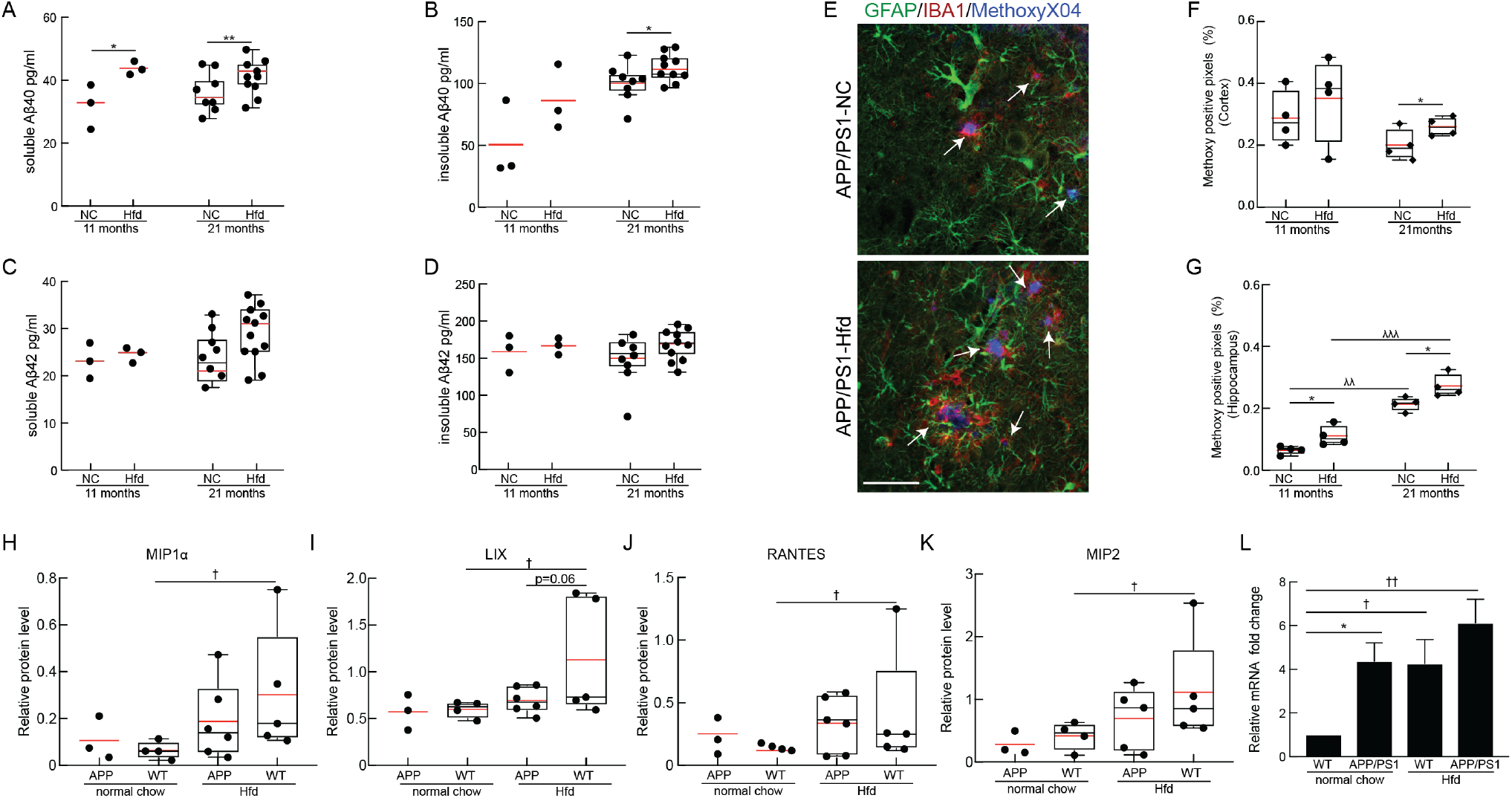
Immunohistochemical and molecular analysis of brain tissue of APP/PS1 and WT mice fed a Hfd or normal chow. PBS-soluble fraction of Aß40 (A) and Aß42 (C) monomeric species and formic acid-soluble fraction of Aß40 (B) and Aß42 (D) monomeric species, measured using ELISA from the brains of APP/PS1 mice on Hfd (started at 4 months of age) or normal chow for mice at 11 and 21 months of age. Animal numbers for all measurements — 11 months: APP/PS1-Hfd: n=3; APP/PS1-NC: n=3; 21 months: APP/PS1-Hfd: n=10; APP/PS1-NC: n=9); *p<0.05 and **p<0.01; Student’s t-test. (E) Representative confocal images of cortical immunostained sections from 21-month old APP/PS1-NC mice on normal chow (top) or Hfd (bottom), stained with MethoxyX04 for amyloid plaques (blue, plaques indicated with white arrows), anti-IBA1 for microglia (red), and anti-GFAP for astrocytes (green). Scale bar = 50μm. Percentage of pixels above threshold for Methoxy-X04 in the cortex (F) and hippocampus (G) for 11 and 21 month old APP/PS1 mice on normal chow and Hfd. Animal numbers for all measurements — 11 months: APP/PS1-NC: n=3; WT-NC: n=3; APP/PS1-Hfd: n=3; WT-Hfd: n=4; 21 months: APP/PS1-NC: n=4; WT-NC: n=4; APP/PS1-Hfd: n=4; WT-Hfd: n=3; * p<0.05 between genotypes (APP/PS1 vs. WT); ^λ^ p<0.05, ^λ λ^ p<0.01, and ^λ λ λ^ p<0.001 between animal ages; Student’s t-test. Relative protein levels of (H) Macrophage inflammatory protein 1 alpha (MIP1α) (I) CXCL5 (LIX) (J) CCL5 (RANTES) and (K) CXCL2 (MIP2) for APP/PS1 and WT mice on normal chow or Hfd at 21 months of age. Animal numbers for all measurements — APP/PS1-NC: n=3; WT-NC: n=4; APP/PS1-Hfd: n=5; WT-Hfd: n=5; † p<0.05 between diets (Hfd vs. NC); one-way ANOVA with post-hoc pair-wise comparisons using Dunn’s multiple comparison correction. (L) Expression of endogenous murine APP mRNA from brain tissue of APP/PS1 and WT mice on a Hfd or normal chow at 21 months of age, normalized to GAPDH mRNA levels. Animal numbers for all measurements — APP/PS1-NC: n=4; WT-NC: n=3; APP/PS1-Hfd: n=4; WT-Hfd: n=4); * p<0.05 between genotypes (APP/PS1 vs. WT); † p<0.05, †† p<0.01 between diets (Hfd vs. NC); Student’s t-test.

To assess brain inflammation, we stained for astrocytes (GFAP) and microglia (Iba1) in serial sections from cortex and hippocampus from mice from all four groups sacrificed at 11 and 21 months of age (Fig. 2E and Supplementary Fig. 1A). For animals on normal chow, we found increased microglia density in APP/PS1 animals, as compared to WT mice, with this difference most clear in the hippocampus (Supplementary Fig. 1B and C). Additionally, we found that the high fat diet increased microglia density in both WT and APP/PS1 mice at 11 months of age, with this increase, again, most clear in the hippocampus (Supplementary Fig. 1B and C). By 21 months of age, the density of microglia had increased in the WT-NC group, dampening the differences between APP/PS1 and WT and between Hfd and NC groups (Supplementary Fig. 1B and C). We did not observe notable differences in astrocyte density between any group at any age (Supplementary Fig. 1D and E).

To further explore the brain inflammation due to APP/PS1 genotype and high fat diet, we performed a cytokine array on all four groups of mice sacrificed at 21 months of age. Overall, we did not notice large changes in expression across a broad range of cytokines between WT-NC and APP/PS1-NC, but we did observe increases in several cytokines in the high fat diet groups (Supplementary Figs. 2 and 3). Interestingly, the largest changes were in the WT-Hfd diet group compared to WT-NC animals, with a more modest effect from the high fat diet evident in the APP/PS1 mice. In particular, several genes involved in leukocyte adhesion and recruitment were elevated in WT-Hfd mice, including the macrophage inflammatory protein 1 alpha (MIP1α), CXCL5 (Lix), RANTES/CCL5, and CXCL2 (MIP2) (Fig. 2H-K and Supplementary Figs. 2 and 3). Finally, we used quantitative PCR to measure the transcript level of the mouse endogenous APP gene and found an increase in WT-Hfd as compared to WT-NC mice. APP/PS1-NC animals showed a similar level of endogenous APP gene expression as the WT-Hfd, with APP/PS1-Hfd trending toward even higher expression. Taken together, this data indicates that Hfd diet can induce the mouse APP promotor (Fig. 2L).

As these four cohorts of animals aged, we periodically analyzed the mice for alterations in short-term memory using the Y-maze and object replacement (OR) tests. At 8 months of age, there were no differences in spontaneous alternation between groups in the Y-maze, a measure of working memory (Fig. 3A and Supplementary Fig. 4A). While WT-NC group displayed the expected short-term memory performance in the OR task (~60% of object exploration time focused on the moved object), there was a trend toward impaired performance in all other groups, with the exploration time of the moved object trending toward chance (Fig. 3C; Supplementary Fig. 4I). By 10 months of age, there was a clear impairment in the spontaneous alternation score for APP/PS1 as compared to WT mice, irrespective of diet (Fig. 3A and Supplementary Fig. 4B). For mice eating normal chow, there was also an impairment of the APP/PS1 animals relative to WT mice on the OR preference score (Fig. 3C and Supplementary Fig. 4J). The WT-Hfd group also showed a trend toward impaired performance on the OR task. These groupwise differences were maintained at 19 months of age, with the exception of an aberrant, modest performance improvement on the Y-maze task for the APP-NC group (Fig. 3A and C, and Supplementary Fig. 4C and K). The number of arm entries in the Y-maze task (Fig. 3B and Supplementary Fig. 4E-H) and the total object exploration time in the OR task (Fig. 3D and Supplementary Fig. 4M-P) did not differ significantly between any groups across all ages.

**Figure 3:**
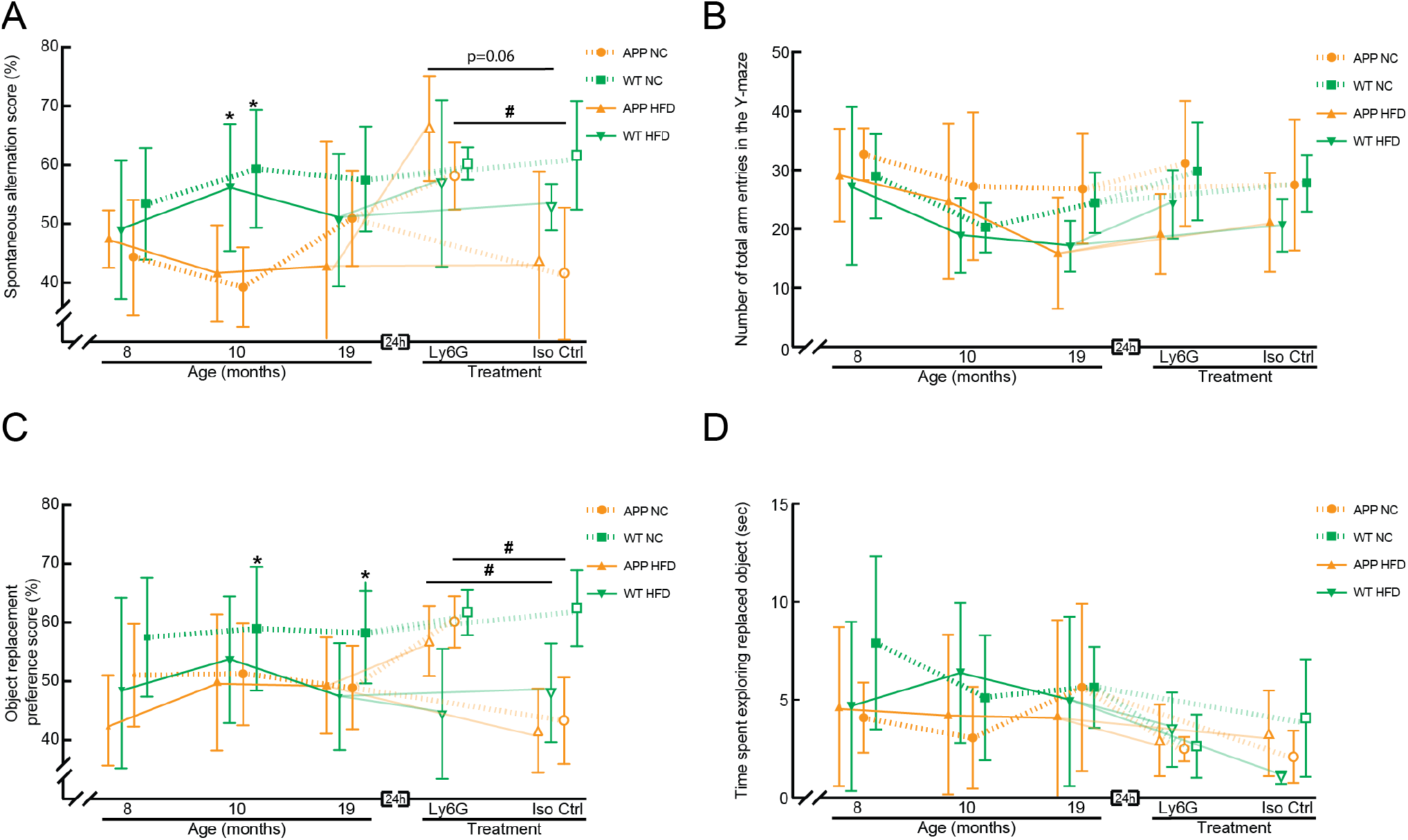
Behavioral assays for short-term memory in APP/PS1 and WT mice on a Hfd or normal chow as a function of age and after treatment with α-Ly6G or Iso-Ctr antibodies. (A) Groupwise averages of spontaneous alternation score and (B) number of arm entries in the Y-maze taken at baseline for mice at 8, 10, and 19 months of age, and taken 24 hrs after α-Ly6G or Iso-Ctr antibody administration at the 19 month time point (4 mg/kg animal weight, intraperitoneal). (C) Groupwise averages of object replacement preference score and (B) time spent at the replaced object in the object replacement task taken at baseline for mice at 8, 10, and 19 months of age, and taken 24 hrs after α-Ly6G or Iso-Ctr antibody administration at the 19 month time point (4 mg/kg animal weight, intraperitoneal). Animal numbers for all measurements — 8 and 10 months: APP/PS1-NC: n=6; WT-NC: n=9; APP/PS1-Hfd: n=10; WT-Hfd: n=11; 19 months: APP/PS1-NC α-Ly6G: n=5; APP/PS1-NC Iso-Ctr n=4; WT-NC α-Ly6G: 7; WT-NC Iso-Ctr: 6; APP/PS1-Hfd α-Ly6G: n=5; APP/PS1-Hfd Iso-Ctr: n=5; WT-Hfd α-Ly6G: n=4; WT-Hfd Iso-Ctr: n=5. * p<0.05 between genotypes (APP/PS1 vs. WT); # p<0.05 between treatment groups (Ly6G vs. Iso-Ctr); Kruskal-Wallis one-way ANOVA with post-hoc pair-wise comparisons using Dunn’s multiple comparison test.

We further characterized these groups of mice for differences in social interaction and preference for social novelty using the three-chamber sociability and social novelty test. Briefly, in the first round, we evaluated the preference of a mouse for an empty chamber vs. direct interaction with (or being in the same chamber as) a same-sex, same-age stranger. In the second round, we compared the preference for the first stranger vs. a new, novel stranger. We found that all animal groups showed similar preferences for social interaction with a stranger and similar preferences for a novel stranger at 10 months of age (Fig. 4 and Supplementary Fig. 5). At 19 months of age, the APP/PS1-Hfd, APP/PS1-NC, and WT-Hfd mice showed decreased preference for social interaction with a stranger and decreased preference for a novel stranger, as compared to the WT-NC mice, with these differences clearest when evaluating time spent in the chamber rather than direct social interaction (Fig. 4B and D, and Supplementary Fig. 5).

**Figure 4:**
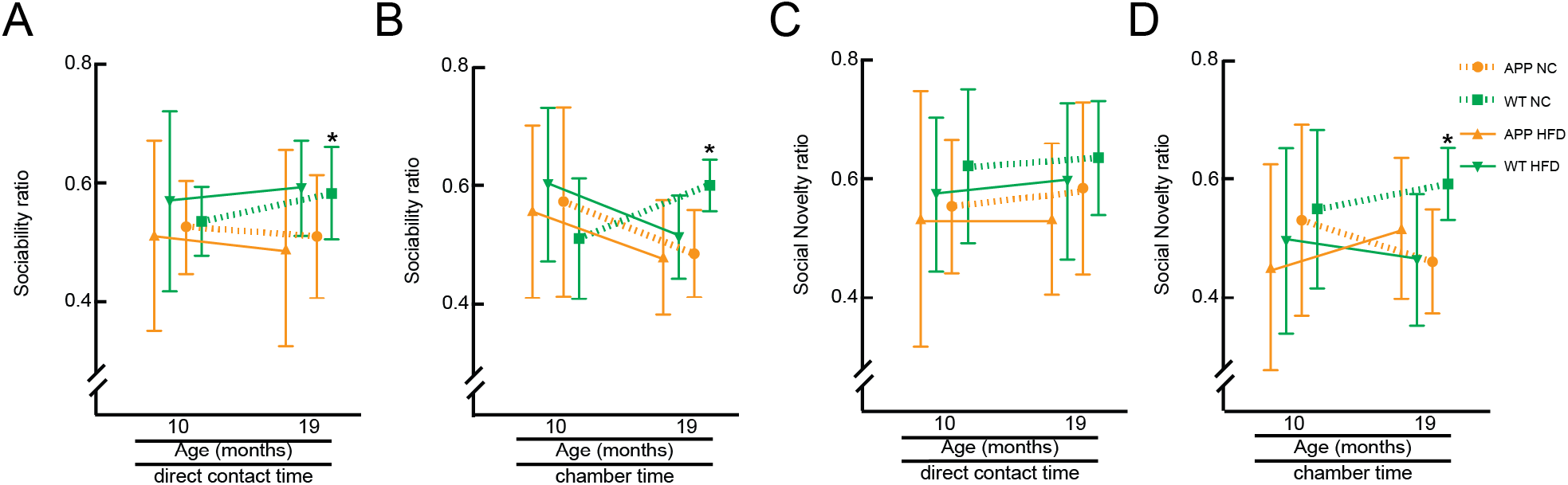
Behavioral assays for social interaction and novelty in APP/PS1 and WT mice on a Hfd or normal chow at 10 and 19 months of age. Groupwise averages for the 3-chamber sociability score, measured as direct interaction time with the stranger vs. an empty chamber (A) and time spent in chamber with the stranger vs. the empty chamber (B), at 10 and 19 months of age. Groupwise averages for the 3-chamber social novelty score, measured as direct interaction time with the novel stranger vs. the old stranger (C) and time spent in chamber with the novel stranger vs. the chamber with the old stranger (D), at 10 and 19 months of age. Animal numbers for all measurements — 10 months: APP/PS1-NC: n=9; WT-NC: n=11; APP/PS1-Hfd: n=13; WT-Hfd: n=13; 19 months: APP/PS1-NC: n=5; WT-NC: n=8; APP/PS1-Hfd: n=10; WT-Hfd: n=11; *p<0.05 between genotypes (APP/PS1 vs. WT); one-way ANOVA with Holm-Šídák multiple comparisons correction.

We also assessed sensory-motor function using a balance beam walk, where the time to cross the beam and the number of hind paw slips were quantified for a 6-mm diameter, 80-cm long beam. For animals on the normal chow, of both APP/PS1 and WT genotypes, we found similar ~10-s crossing times and only a few hind paw slips for mice at both 10 and 19 months of age (we note, however, the aberrantly poor performance of the WT-NC group at 10 months of age) (Fig. 5A and B, Supplementary Fig. 6A, B, D, and E). WT mice on the Hfd showed performance comparable to animals on normal chow at 10 months of age, but trended toward poorer performance by 19 months of age. While APP/PS1 phenotype or Hfd alone did not strongly impact balance beam performance, the APP/PS1 mice on the Hfd showed a synergistic impact of genotype and diet, with significantly slower crossing times and increased number of hind paw slips at both 10 and 19 months of age, as compared to APP/PS1 on normal chow or WT mice on the Hfd (Fig. 5A and B, Supplementary Fig. 6A, B, D, and E).

**Figure 5:**
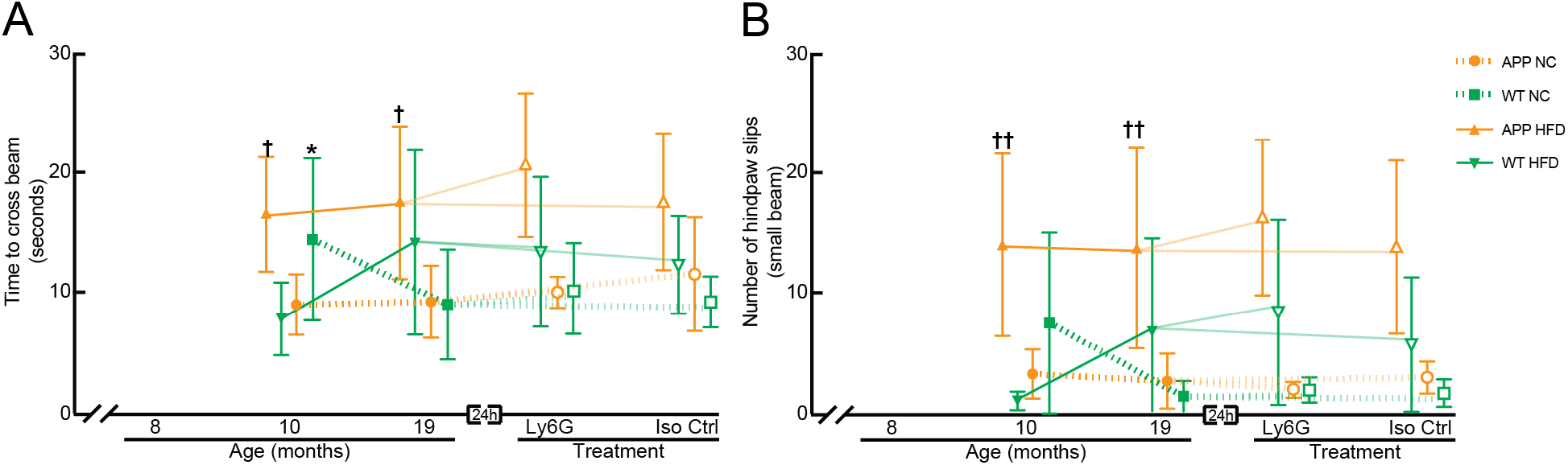
Behavioral assays for sensory-motor function in APP/PS1 and WT mice on a Hfd or normal chow as a function of age and after treatment with α-Ly6G or Iso-Ctr antibodies. Groupwise averages of balance beam performance measured by (A) the time to cross the beam and (B) the number of hindpaw slips while crossing for mice at 10 and 19 months of age, and taken 24 hrs after α-Ly6G or Iso-Ctr antibody administration at the 19 month time point (4 mg/kg animal weight, intraperitoneal). Animal numbers for all measurements — 10 months: APP/PS1-NC: n=6; WT-NC: n=9; APP/PS1-Hfd: n=10; WT-Hfd: n=11; 19-months: APP/PS1-NC α-Ly6G: n=5; APP/PS1-NC Iso-Ctr n=4; WT-NC α-Ly6G: n=7; WT-NC Iso-Ctr: n=6; APP/PS1-Hfd α-Ly6G: n=5, APP/PS1-Hfd Iso-Ctr: n=5; WT-Hfd α-Ly6G: n=4; WT-Hfd Iso-Ctr: n=5. * p<0.05 between genotypes (APP/PS1 vs. WT); † p<0.05, †† p<0.01 between diets (Hfd vs. NC); one-way ANOVA with post-hoc pair-wise comparisons using Dunn’s multiple comparison test; mixed-model to determine synergistic effect of genotype and diet - (time to cross: p = 0.06; number of hindpaw slips: p = 0.008).

On the whole, the behavioral, histopathological, and molecular data above suggest that the APP/PS1 mice on the high fat diet in this study showed a similar exacerbation of Alzheimer’s-related effects as in previous studies[16, 17, 22]. We next sought to determine the role of capillary stalling and CBF deficits in this Hfd-linked exacerbation of Alzheimer’s related symptoms. Based on our previous findings that administration of antibodies against Ly6G (α-Ly6G) led to improvements in performance on Y-maze and OR tasks in APP/PS1 mice within hours and persisting for at least a day [32, 33], we assessed whether α-Ly6G administration improved memory function in the 19 month old mice, across all groups. For APP/PS1-NC and APP/PS1-Hfd mice, treatment with α-Ly6G led to more spontaneous alternation in the Y-maze (Fig. 3A and Supplementary Fig. 4D) and higher preference scores for the moved object in the OR task (Fig. 3C and Supplementary Fig. 4L) at 24 hours after treatment, as compared to the 19 month baseline data. In contrast, treatment with α-Ly6G did not change the performance of the WT-NC and WT-Hfd mice on either Y-maze or OR tasks, so the impairment of WT-Hfd mice on the OR task was retained after α-Ly6G treatment. Treatment with Iso-Ctr antibodies did not impact performance of any group for either task. The number of arm entries in the Y-maze (Fig. 3B and Supplementary Fig. 4H) and the total object exploration time in the OR task (Fig. 3D and Supplementary Fig. 4P) was not impacted by α-Ly6G or Iso-Ctr treatment for any group. Treatment with α-Ly6G or Iso-Ctr antibodies did not lead to any changes in crossing times or number of slips for any groups in the balance beam assay, relative to the 19-month baseline data (Fig. 5A and B, Supplementary Fig. 6C and F).

We next sought to determine if mice eating the high fat diet exhibited increased capillary stalling and reduced CBF. Image stacks of the cortical capillary bed were taken using 2PEF microscopy across all four groups of mice (Fig. 6A-D) and ~96,000 individual capillary segments were scored as flowing or stalled (Fig. 6E). Consistent with our previous reports [32, 33], we found an increase in the fraction of capillaries with stalled blood flow in APP/PS1 mice as compared to WT, regardless of diet (percentage of capillaries stalled at 11 months (21 months) of age: APP/PS1-NC: 2% (1.1%); APP/PS1-Hfd: 1.7% (1.3%); WT-NC: 0.6% (0.3%); WT-Hfd: 0.5% (0.5%)). Consistent with our previous findings, these data show a trend toward a decrease in the incidence of stalled capillaries with increasing age in the APP/PS1 mice. We observed no impact of diet on the incidence of stalled capillaries in either APP/PS1 or WT mice (Fig. 6F and Supplementary Fig. 8A). We distinguished different potential causes of stalled capillaries using an *in vivo* labeling strategy where leukocytes are labeled by both Rhodamine 6G (mitochondria) and Hoechst (DNA), platelets are labeled by Rhodamine 6G only, and RBCs are unlabeled and appear as dark patches in the fluorescently-labeled blood plasma (Texas Red-dextran)[32, 40]. In the APP/PS1-NC mice, about 86% of the capillary stalls had a leukocyte in the stalled capillary segment, while 5% had platelets, and 9% had only RBCs. WT-NC mice not only had a lower total incidence of stalled capillaries, they also had different cellular compositions, with 0% containing leukocytes, 50% platelets, and 50% only RBCs. For mice on the Hfd, APP/PS1 animals had similar ratios to the normal chow animals (80% leukocytes, 15% platelets, and 5% only RBCs), while WT-Hfd mice exhibited a shift towards more stalls being caused by leukocytes (60% leukocytes, 0% platelets, and 40% only RBCs).

**Figure 6:**
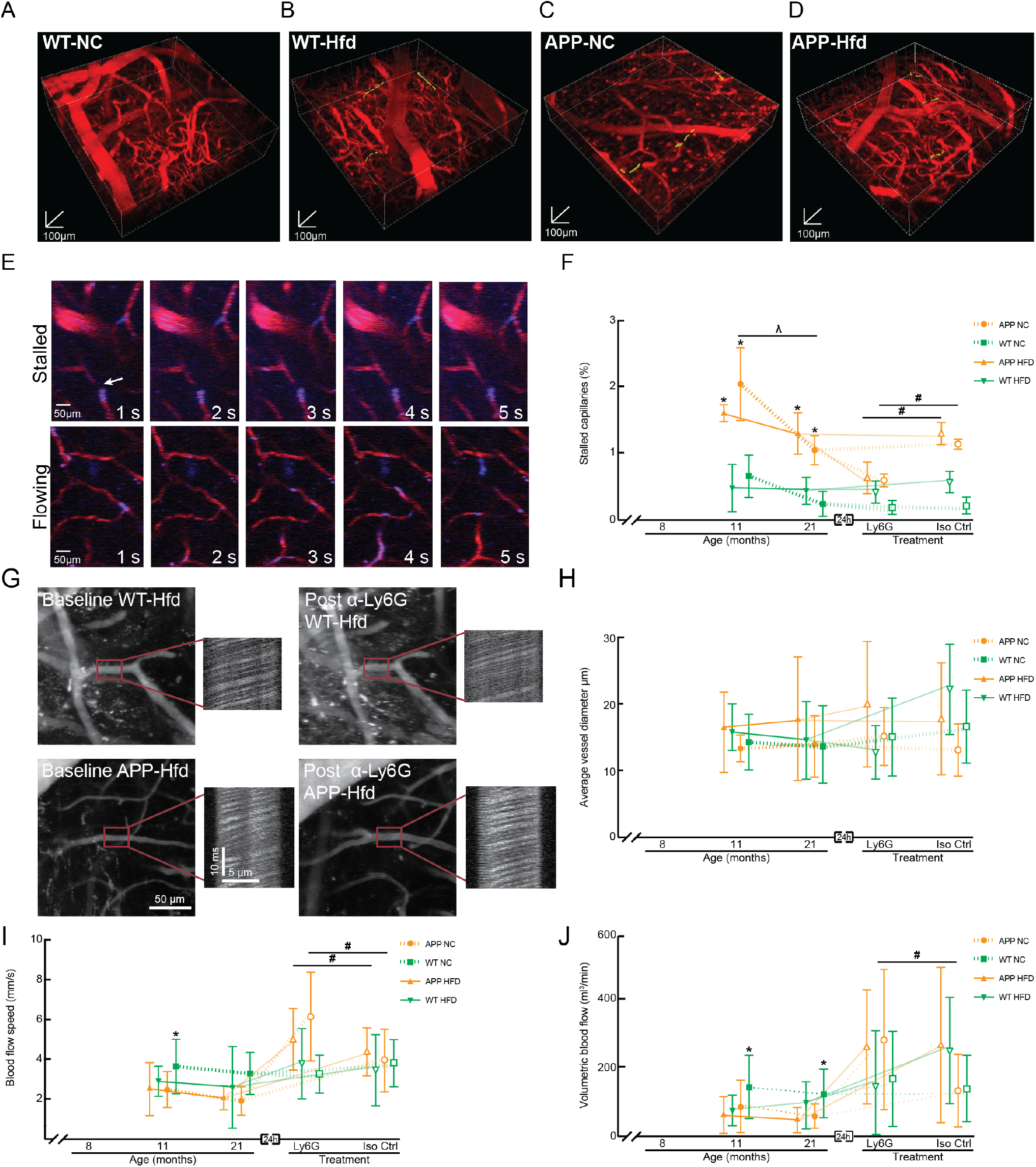
Incidence of non-flowing capillaries and ateriole blood flow speeds in APP/PS1 and WT animals on a Hfd or normal chow. (A-D) Renderings of 2PEF image stacks of cortical vasculature (red; Texas Red dextran) with stalled capillaries indicated by yellow highlights for APP/PS1 and WT mice on Hfd or NC. 3D scale bars are 100 μm. (E) Sequence of 2PEF images used to score individual capillaries as flowing or stalled based on the motion of unlabeled blood cells (dark patches in vessel lumen) within the fluorescently labeled blood plasma (red). Rhodamine 6G (blue) labels leukocytes and platelets. Scale bar represents 50 μm. (F) Groupwise averages of the fraction of cortical capillaries with stalled blood flow at baseline for mice at 11 and 21 months of age and taken 24 hours after α-Ly6G or Iso-Ctr antibody administration at the 21 month time point (4 mg/kg animal weight, intraperitoneal). Animal numbers for all measurements — 11 and 21 months: APP/PS1-NC: n=3; WT-NC: n=3; APP/PS1-Hfd: n=3; WT-Hfd: n=3; 21 months: APP/PS1-NC α-Ly6G: n=3; APP/PS1-NC Iso-Ctr n=3; WT-NC α-Ly6G: 3; WT-NC Iso-Ctr: 3; APP/PS1-Hfd α-Ly6G: n=3; APP/PS1-Hfd Iso-Ctr: n=3; WT-Hfd α-Ly6G: n=3; WT-Hfd Iso-Ctr: n=3. * p<0.05 between genotypes (APP/PS1 vs. WT); # p<0.05 between treatment groups (Ly6G vs. Iso-Ctr); Kruskal-Wallis one-way ANOVA with post-hoc pair-wise comparisons using Dunn’s multiple comparison test. (G) Representative 2PEF images and line scan data from penetrating arterioles before and after α-Ly6G injection from WT-Hfd (upper panels) and APP-Hfd (lower panels). Scale bar represents 50 μm. Groupwise averages of average vessel dimeter (H), RBC flow speed (I), and volumetric blood flow (J) taken at baseline for mice at 11 and 21 months of age, and taken 24 hrs after α-Ly6G or Iso-Ctr antibody administration at the 21 month time point (4 mg/kg animal weight, intraperitoneal). Penetrating ateriole numbers for all measurements — 11 and 21 months: APP/PS1-NC: n=26; WT-NC: n=37; APP/PS1-Hfd: n=27; WT-Hfd: n=26; 21 months: APP/PS1-NC α-Ly6G: n=21; APP/PS1-NC Iso-Ctr n=12; WT-NC α-Ly6G: 13; WT-NC Iso-Ctr:9; APP/PS1-Hfd α-Ly6G: n=18; APP/PS1-Hfd Iso-Ctr: n=24; WT-Hfd α-Ly6G: n=15; WT-Hfd Iso-Ctr: n=11. * p<0.05 between genotypes (APP/PS1 vs. WT); # p<0.05 between treatment groups (Ly6G vs. Iso-Ctr); ^λ^ p<0.05 between animal ages; Kruskal-Wallis one-way ANOVA with post-hoc pair-wise comparisons using Dunn’s multiple comparison test.

We previously showed in multiple mouse models of APP overexpression (including APP/PS1 mice) that neutrophils were the leukocyte responsible for nearly all the capillary stalls and that an antibody against the neutrophil-specific surface protein Ly6G (α-Ly6G) rapidly decreased the incidence of stalled capillaries [32]. We therefore determined if a single dose of α-Ly6G reduced the number of capillary stalls in the mice studied here. In APP/PS1-NC and APP/PS1-Hfd mice, α-Ly6G treatment reduced the incidence of stalled capillaries to near WT levels within one hour, while treatment with isotype control antibodies had no impact. In WT-NC and WT-Hfd mice, administration of either α-Ly6G or isotype control antibodies did not alter the already low incidence of stalled capillaries (Fig. 6F and Supplementary Fig. 7B).

To assess the impact of the high fat diet on cortical blood flow in APP/PS1 and WT mice, we measured vessel diameter and centerline red blood cell flow speed (using line scans [40]) in 5-15 penetrating arterioles in each mouse at baseline and about one hour after treatment with α-Ly6G or isotype control antibodies (Fig. 6G). The average diameter of the penetrating arterioles measured were similar across all four groups at both ages and were not impacted by treatment with α-Ly6G or isotype control antibodies (Fig. 6H and Supplementary Fig. 7E and H). Median blood flow speeds were reduced by ~40% (20%) in APP/PS1-NC mice at 11 (21) months of age, relative to the WT-NC animals (Fig. 6I and Supplementary Fig. 7C and I). Treatment with α-Ly6G antibodies increased penetrating arteriole speed by 34% in the 21 month old APP/PS1-NC mice, while isotype control antibodies did not increase flow speeds (Fig. 6I and Supplementary Fig. 7F). At 11 months of age, the APP/PS1-Hfd mice trended toward a ~30% reduction in median penetrating arteriole blood flow speeds relative to the WT-Hfd mice, although the flow speeds measured in the APP/PS1-Hfd mice showed unusual heterogeneity (Fig. 6I and Supplementary Fig. 7C and J). By 21 months of age, there was no difference in flow speeds between APP/PS1-Hfd and WT-Hfd mice. Treatment with α-Ly6G antibodies increased flow speeds by 14% in APP/PS1-Hfd mice at 21 months of age, while isotype control antibodies did not (Fig. 6I and Supplementary Fig. 7F). The high fat diet did not lead to notable differences in flow speed in either WT or APP/PS1 mice, as compared to normal chow. The trends described above for flow speed were similar for volumetric blood flow, which is calculated from the centerline flow speed and vessel diameter (Fig. 6J and Supplementary Figs. 7D and G).

## Discussion

In this study we tested the idea that a synergistic increase in the number of non-flowing capillaries could be a mechanistic link between obesity and increased severity and incidence of AD. To do this, we fed mouse models of AD a Hfd and explored the impact on short-term memory function, sociability, sensory-motor function, amyloid deposition, brain inflammation, capillary stalling, and cerebral blood flow.

Several studies have reported that feeding mouse models of APP overexpression a high fat diet leads to more severe short-term memory deficits [17, 22, 44]. In our work, we did not observe poorer short term memory function in APP/PS1 mice on a Hfd, as compared to normal chow, perhaps reflecting the sensitivity of the memory tests used here or reflecting differences in the composition, time of onset, duration of administration of the Hfd, or the control diet, as compared to other studies. Indeed, there are highly variable results reported on the effect of a western Hfd on memory function in AD mouse models [19]. Consistent with previous studies [16, 17], WT mice on the Hfd tended to show poorer memory function than WT mice on normal chow in our experiments. Treatment with α-Ly6G to reduce capillary stalling restored short-term memory function in the APP/PS1 mice on either diet but did not alter the trend toward poorer memory function in the WT-Hfd mice, suggesting other mechanisms underlie the trend toward short-term memory deficits in these mice. Both APP/PS1 genotype and eating a Hfd was associated with a tendency for impaired social interactions, as compared to WT mice on normal chow. In contrast, eating a Hfd was associated with increased social interaction in an AD mouse model that replaces the endogenous APP gene with mutant, human APP [19], perhaps reflecting differences between APP overexpression and APP knock-in AD mouse models [45]. Both patients and mouse models of AD exhibit sensory-motor deficits including gait disturbances, bradykinesia, and reduced overall activity levels [46, 47]. In the balance beam test of sensory-motor function, we did not observe a deficit in APP/PS1 mice on a normal diet and saw a weak trend toward impairment in WT mice on the Hfd, but we did find a strong impairment with the Hfd in the APP/PS1 mice. Treatment with α-Ly6G did not improve this synergistic deficit in sensory-motor function.

The Hfd was associate with an increased concentration of some species of amyloid-beta and an increased density of amyloid plaques, increased activation of microglia, and increased expression of inflammatory cytokines in the APP/PS1 mice. In WT animals, the Hfd led to similar increases in inflammatory cytokine expression as in APP/PS1 mice. Previous studies have also suggested increased amyloid load, increased monomeric Aß species, increased hyperphosphorylated tau, and more severe inflammation in AD mouse models fed a high fat diet [17, 48–51], although some studies have reported different trends [19, 52], reflecting the variability of the impact of a high fat diet or differences in experimental design. Recent work has shown minimal exacerbation of AD-like pathology with a Hfd in the App^NL/NL^ knock-in mouse model of AD, suggesting that the overexpression of mutant APP, as in most AD mouse models, may also influence the impact of a Hfd [19, 45, 53]. Interestingly in APP/PS1 mice, the worsened pathology with a high fat diet could be reversed by changing back to a low-fat control diet [18]. In our experiments, the expression of endogenous murine APP was increased with the Hfd in WT mice, as well as in APP/PS1 mice on normal chow (with a trend toward a further synergistic increase with Hfd), suggesting that Hfd can induce gene expression programs that may contribute to an Alzheimer’s like phenotype. Similar increases in the expression of endogenous murine APP has been also reported in a mouse model of Down syndrome [54, 55]. In summary, the APP/PS1 mice fed a Hfd in this study broadly reproduced the phenotype of impaired behavior and increased brain inflammation that previous research on obesity and AD in mouse models has reported.

We did not observe an increase in the number of non-flowing capillaries or a larger CBF deficit in the APP/PS1 mice fed a Hfd. Independent of whether the APP/PS1 mice were on a Hfd or normal chow, reducing capillary stalls reliably led to improved memory performance and increased CBF. Taken together, this data suggests that there is not an increase in the number of non-flowing capillaries due to a high fat diet and that such diets exacerbate AD pathology and symptoms through other mechanisms.

While none of these trends rose to significance, we do note that by 21 months of age, there was a doubling of the number of stalled capillaries in WT mice on the Hfd and a shift toward these non-flowing capillaries to be associated with leukocytes. There was also a decrease in the difference in CBF between APP/PS1 and WT mice on the Hfd, as compared to normal chow. Treatment with α-Ly6G did not improve CBF (nor did it rescue any other phenotypes) in WT-Hfd mice. These trends suggest there may be a weak effect that increases non-flowing capillaries, which may not involve neutrophils, and there are certainly other mechanisms contributing to impaired CBF with Hfd, such as increased tonic vasoconstriction [56], or impaired vasodilation in response to changing physiological conditions, such as hypercapnia, both associated with high fat diet consumption and obesity [28, 29, 57–59].

In summary, a western Hfd induces a complex series of changes in APP/PS1 mice, including an increased inflammatory response, increased concentration of amyloid-beta species, and exacerbated behavioral deficits, especially in sensory-motor function. However, we did not observe a synergistic increase in capillary stalling or CBF deficit in APP/PS1 mice on the Hfd, suggesting that other mechanisms underlie the synergistic impact of a Hfd and APP/PS1 genotype. We did show that blocking neutrophil adhesion improved CBF and short-term memory function in APP/PS1 mice on the Hfd, suggesting that improving blood flow is still possible and still improves brain function in combined obesity and AD.

## Acknowledgements

Confocal imaging data was acquired through the Cornell University Biotechnology Resource Center, with NIH funding (grant number RR025502) for the shared Zeiss LSM 710 Confocal.

