## Supplemental 1-7 for "High fat diet worsens pathology and impairment in an Alzheimer’s mouse model, but not by synergistically decreasing cerebral blood flow"

**Supplemental Figures:**

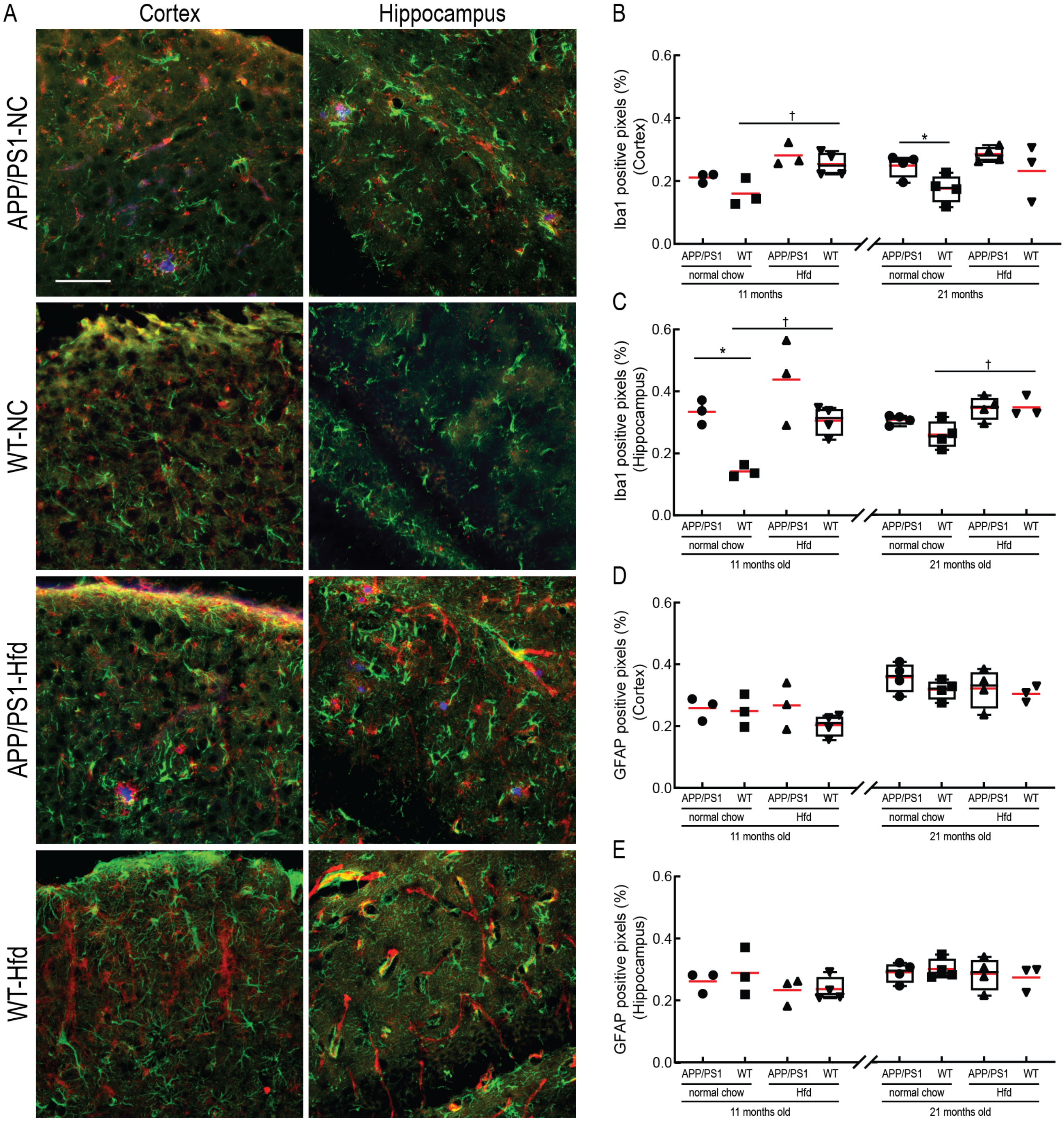


**Supplementary Figure 1: Immunostaining of cortex and hippocampus in APP/PS1 and WT mice on a Hfd or normal chow.** (A) Confocal images of cortical (left) and hippocampal (right) regions in APP/PS1 and WT mice on a normal chow (top) or the Hfd (bottom). Anti-GFAP for astrocytes (green), anti-IBA1 for microglia (red), Methoxy-X04 for amyloid plaques (blue). Scale bar indicates 50 µm. (B-E) Fraction of pixels positive for anti-IBA1 in the cortex (B) and hippocampus (C), and fraction of pixels positive for anti-GFAP in the cortex (D) and hippocampus (E) of APP/PS1 and WT mice on a Hfd or normal chow at 11 and 21 months of age. Animal numbers for all measurements — 11 months: APP/PS1-NC: n=3; WT-NC: n=3; APP/PS1-Hfd: n=3; WT-Hfd: n=4; 21 months: APP/PS1-NC: n=4; WT-NC: n=4; APP/PS1-Hfd: n=4; WT-Hfd: n=3; * p<0.05 between genotypes (APP/PS1 vs. WT); p<0.05, p<0.01 between diets (Hfd vs. NC); one-way ANOVA with post-hoc pair-wise comparisons using Dunn's multiple comparison test.

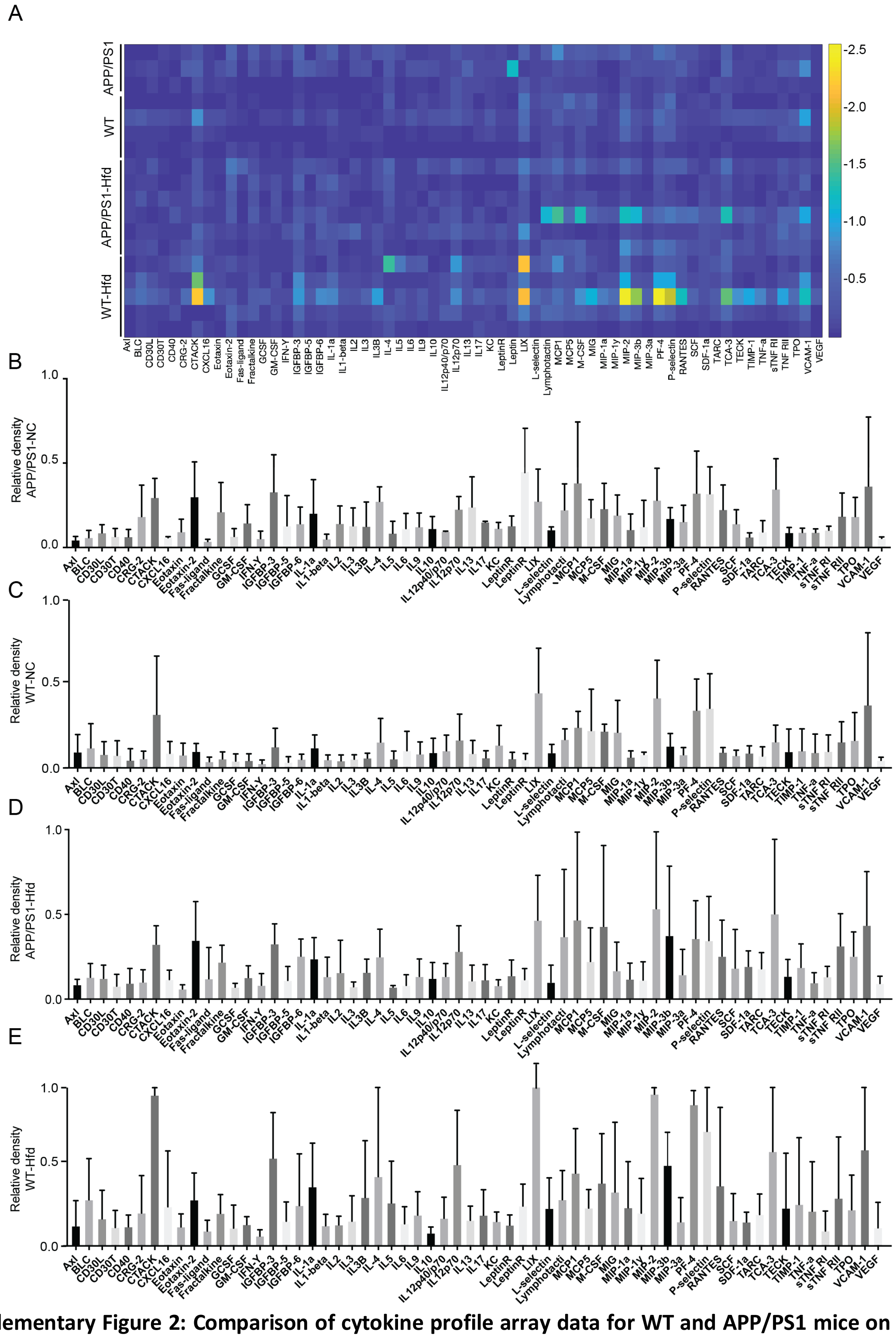


**Supplementary Figure 2: Comparison of cytokine profile array data for WT and APP/PS1 mice on the Hfd or normal chow.** (A) Heatmap of the brain cytokine levels for all WT and APP/PS1 mice on the Hfd or normal chow at 21 months of age. The color bar calibrates the fold changes of expression, relative to manufacturer internal controls. (B-E) Groupwise averaged relative cytokine expression for (B) APP/PS-NC: n=3; (C) WT-NC: n=4; (D) APP/PS1-Hfd: n=6; (E) WT-Hfd: n=5.

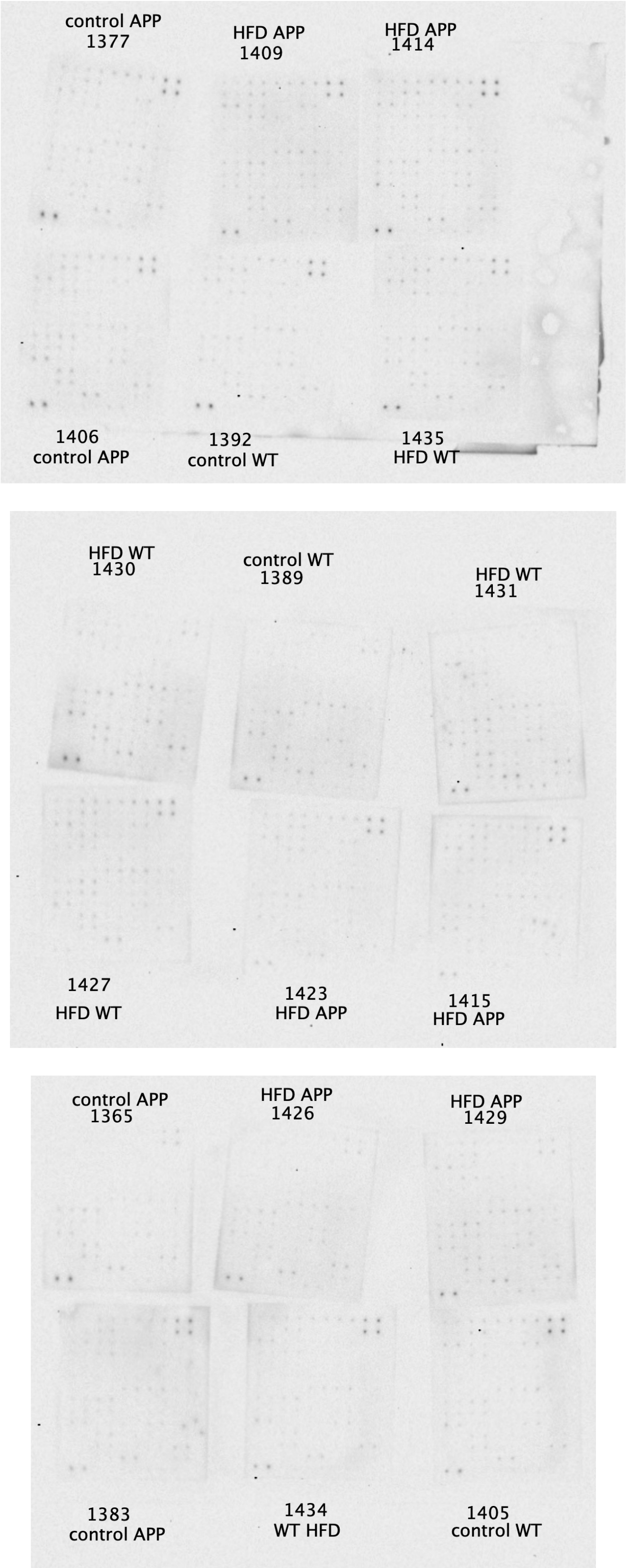


**Supplementary Figure 3: Raw images of cytokine arrays of APP/PS1 and WT mice on a Hfd or normal chow.** Raw images with an exposure time of 2 minutes for each individual membrane. Indicated are the animal numbers and experimental group.

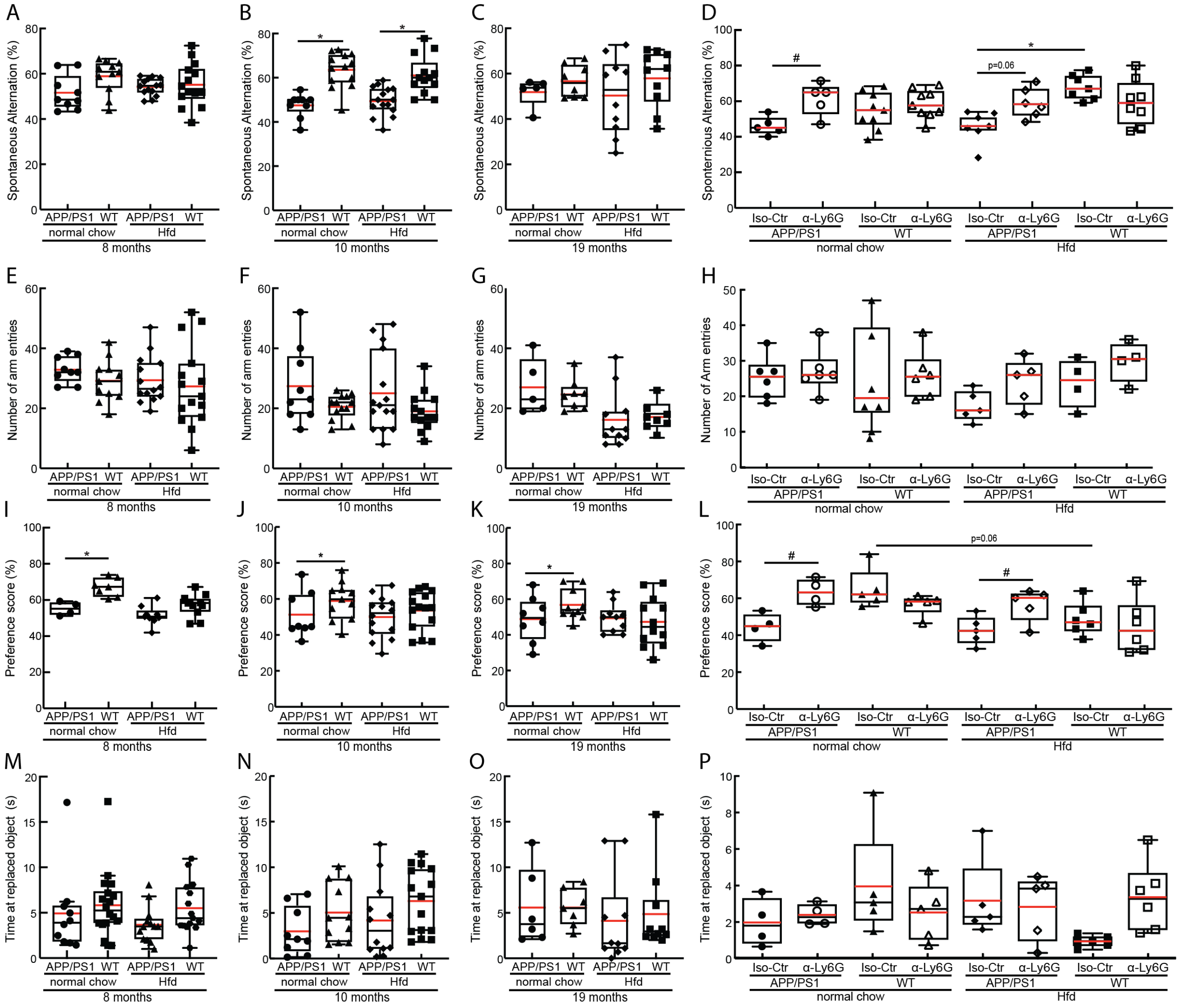


**Supplementary Figure 4: Mouse-by-mouse data for short-term memory tests.** (A-D) Box plots of the spontaneous alternation score and (E-G) number of arm entries from Y-maze experiments, taken at baseline for mice at 8, 10, and 19 months of age, and taken 24 hrs after α-Ly6G or Iso-Ctr antibody administration at the 19 month time point (4 mg/kg animal weight, intraperitoneal). (I-L) Box plots of the preference score and (E-G) number of arm entries from the object replacement task, taken at baseline for mice at 8, 10, and 19 months of age, and taken 24 hrs after α-Ly6G or Iso-Ctr antibody administration at the 19 month time point (4 mg/kg animal weight, intraperitoneal. 8 and 10 months: APP/PS1-NC: n=6; WT-NC: n=9; APP/PS1-Hfd: n=10; WT-Hfd: n=11; 19 months: APP/PS1-NC α-Ly6G: n=5; APP/PS1-NC Iso-Ctr n=4; WT-NC α-Ly6G: 7; WT-NC Iso-Ctr: 6; APP/PS1-Hfd α-Ly6G: n=5; APP/PS1-Hfd Iso-Ctr: n=5; WT-Hfd α-Ly6G: n=4; WT-Hfd Iso-Ctr: n=5. *$p<0.05$ between treatment groups (Ly6G vs. Iso-Ctr); # $p<0.05$ between treatment groups (Ly6G vs. Iso-Ctr), Kruskal-Wallis one-way ANOVA with post-hoc wise comparisons using Dunn's multiple comparison test.

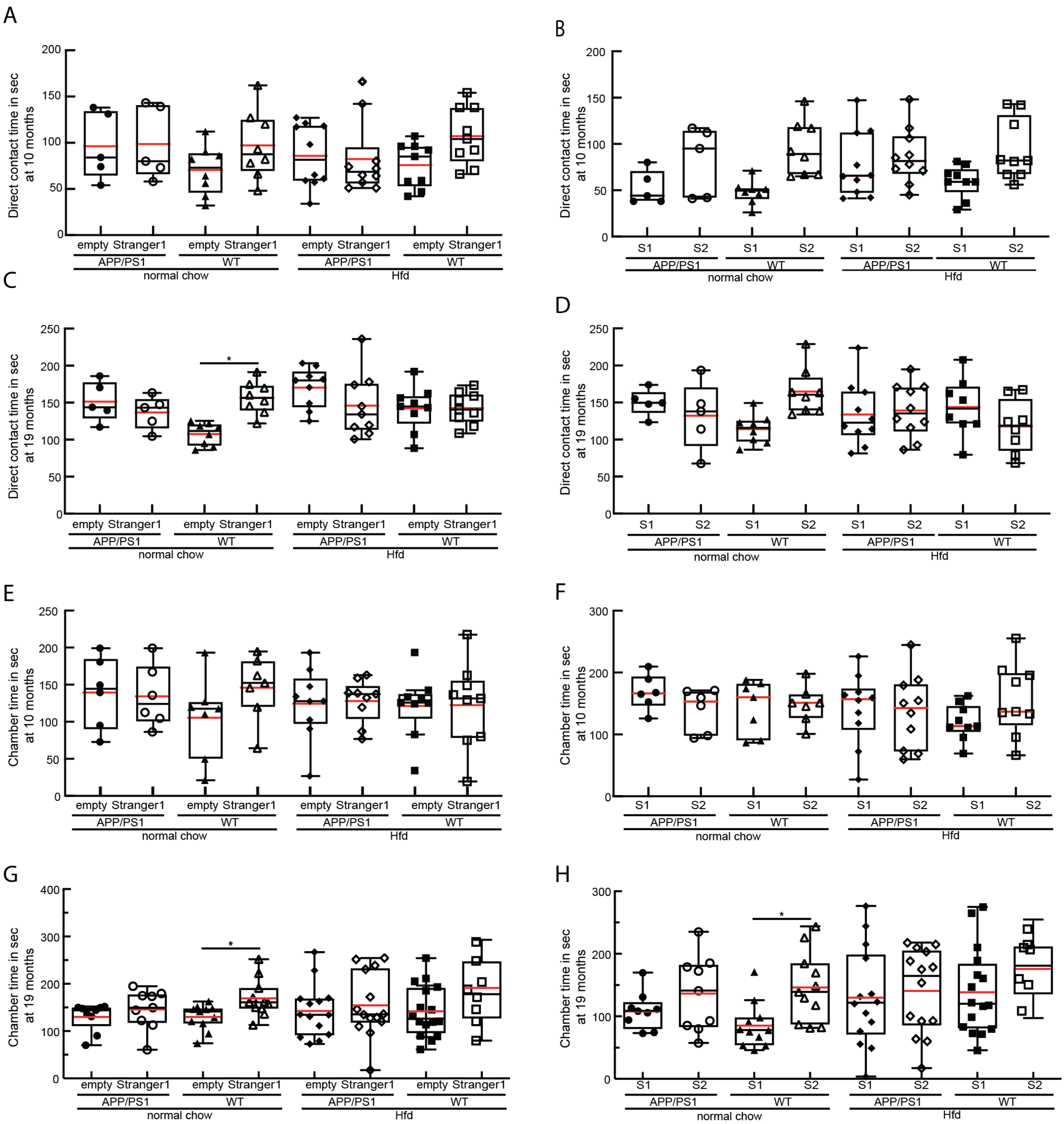


**Supplementary Figure 5: Mouse-by-mouse data for the 3-chamber social interaction and social novelty test.** (A-D) Box plots of direct contact time with Stranger 1 vs the empty chamber in the sociability test (A and C) and with Stranger 2 vs Stranger 1 (B and D) from the social novelty test for APP/PS1 and WT mice on a Hfd or normal chow at 10 months (A and B) or 19 months (C and D) of age. (E-H) Box plots of chamber time with Stranger 1 vs the empty chamber in the sociability test (A and C) and with Stranger 2 vs Stranger 1 (B and D) from the social novelty test for APP/PS1 and WT mice on a Hfd or normal chow at 10 months (A and B) or 19 months (C and D) of age.

Animal numbers for all measurements — 10 months: APP/PS1-NC: n=9; WT-NC: n=11; APP/PS1-Hfd: n=13; WT-Hfd: n=13; 19 months: APP/PS1-NC: n=5; WT-NC: n=8; APP/PS1-Hfd: n=10; WT-Hfd: n=11; *$p<0.05$ and **$p<0.01$; one-way ANOVA with Holm-Šídák multiple comparisons correction.

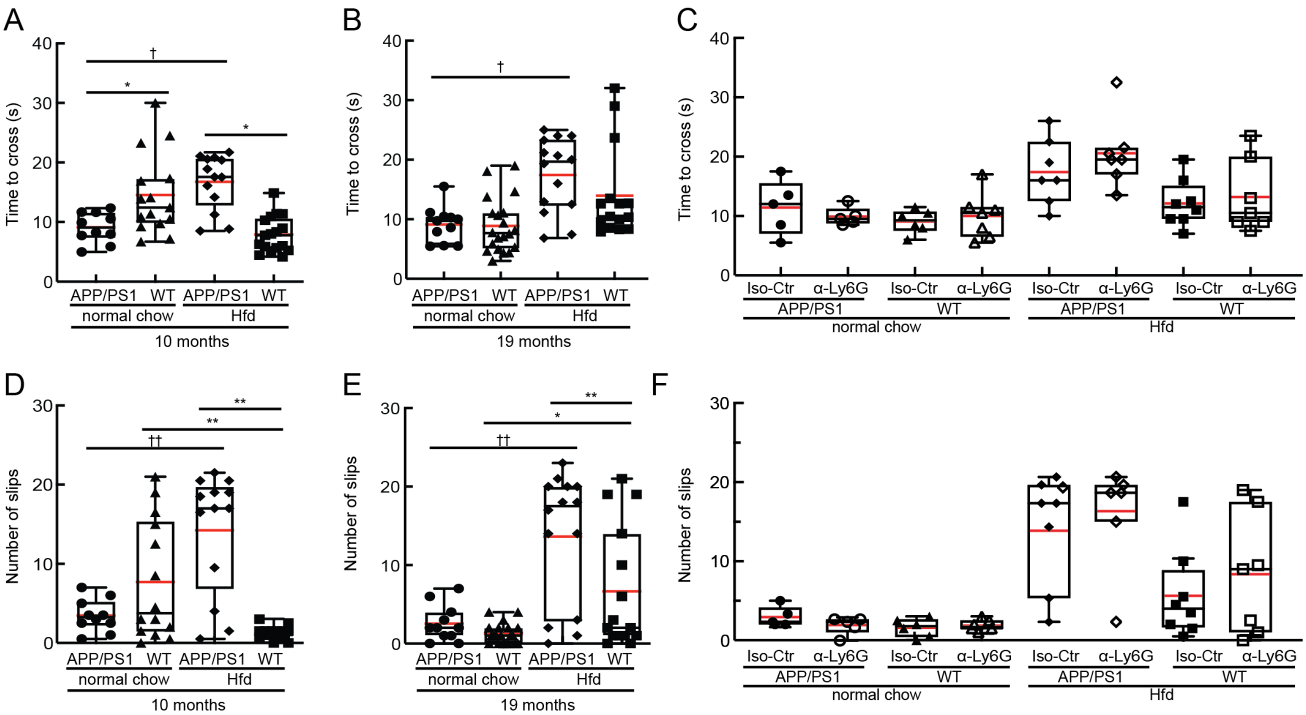


**Supplementary Figure 6: Mouse-by-mouse data for the balance beam walk test of sensory-motor function.** (A-B) Box plots of the time to cross the beam for the balance beam test for mice at 10 and 19 months of age, and (C) time to cross taken 24 hrs after α-Ly6G or Iso-Ctr antibody administration at the 19 month time point (4 mg/kg animal weight, intraperitoneal). (D-E) Box plots of the number of hindpaw slips in APP/PS1 and WT mice on a Hfd or normal chow at 10 and 19 months of age, and (F) number of hindpaw slips taken 24 hrs after α-Ly6G or Iso-Ctr antibody administration at the 19 month time point (4 mg/kg animal weight, intraperitoneal). Animal numbers for all measurements — 10 months: APP/PS1-NC: n=6; WT-NC: n=9; APP/PS1-Hfd: n=10; WT-Hfd: n=11; 19-months: APP/PS1-NC α-Ly6G: n=5; APP/PS1-NC Iso-Ctr n=4; WT-NC α-Ly6G: n=7; WT-NC Iso-Ctr: n=6; APP/PS1-Hfd α-Ly6G: n=5, APP/PS1-Hfd Iso-Ctr: n=5; WT-Hfd α-Ly6G: n=4; WT-Hfd Iso-Ctr: n=5. * $p<0.05$ and ** $p<0.01$ between genotypes (APP/PS1 vs. WT); † $p<0.05$, ††$p<0.01$ between diets (Hfd vs. NC), one-way ANOVA with post-hoc pair-wise comparisons using Dunn's multiple comparison test.

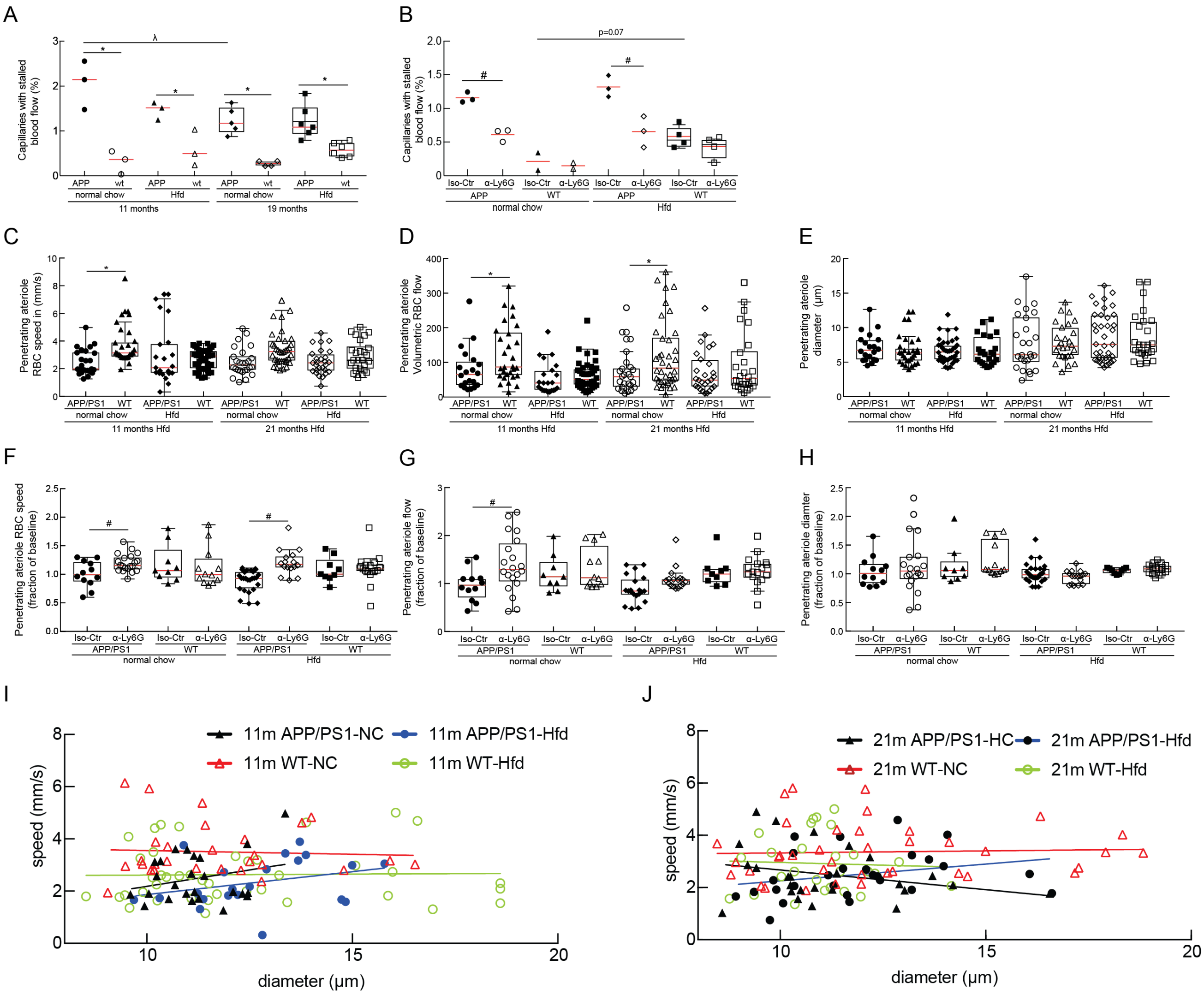


**Supplementary Figure 7: Mouse-by-mouse data for the fraction of capillaries with stalled blood flow and for penetrating arteriole blood flow speeds in APP/PS1 and WT mice on normal chow or Hfd, before and after treatment with α-Ly6G or Iso-Ctr antibodies.** (A) Box plot of the fraction of capillaries with stalled blood flow from 11 and 21 month old APP/PS1 and WT mice on the Hfd or normal chow. Animal numbers for all measurements — 11 month: APP/PS1-NC: n=3; WT-NC: n=3; APP/PS1-Hfd: n=3; WT-Hfd: n=3; 21 months: APP/PS1-NC: n=6; WT-NC: n=6; APP/PS1-Hfd: n=6; WT-Hfd: n=6. * p<0.05 between genotypes (APP/PS1 vs. WT); # p<0.05 between treatment groups (Ly6G vs. Iso-Ctr); λ p<0.05 between animal ages; Kruskal-Wallis one-way ANOVA with post-hoc pair-wise comparisons using Dunn's multiple comparison test. (B) Box plot of the fraction of capillaries with stalled blood flow ~1 hour after α-Ly6G or Iso-Ctr antibody administration (4 mg/kg animal weight, intraperitoneal) for 21 month old mice. Animal numbers for all measurements — APP/PS1-NC α-Ly6G: n=3; APP/PS1-NC Iso-Ctr n=3; WT-NC α-Ly6G: 3; WT-NC Iso-Ctr: 3; APP/PS1-Hfd α-Ly6G: n=3; APP/PS1-Hfd Iso-Ctr: n=3; WT-Hfd α-Ly6G: n=3; WT-Hfd Iso-Ctr: n=3. * p<0.05 for comparisons between genotypes (APP/PS1 vs. WT), Kruskal-Wallis one-way ANOVA with post-hoc pair-wise comparisons using Dunn's multiple comparison test. Box plots of (C) RBC flow speed, (D) volumetric blood flow, and (E) vessel diameter from 11 and 21 month old APP/PS1 and WT mice on the Hfd or normal chow. Penetrating arteriole numbers (mice numbers) for all measurements — 11 months: APP/PS1-NC: n=26 (3); WT-NC: n=37 (3); APP/PS1-Hfd: n=27 (3); WT-Hfd: n=26(3); 21 months: APP/PS1-NC: n=33 (6); WT-NC: n=22 (5); APP/PS1-Hfd: n=42 (7); WT-Hfd: n=26 (6). * p<0.05 between genotypes (APP/PS1 vs. WT), Kruskal-Wallis one-way ANOVA with post-hoc pair-wise comparisons using Dunn's multiple comparison test. (F) RBC flow speed, (G) volumetric blood flow and (H) vessel diameter, all expressed as a fraction of the baseline value, for 21 month old APP/PS1 and WT mice taken 24 hrs after α-Ly6G or isotype control antibody administration. Penetrating arteriole numbers (mice numbers) for all measurements — APP/PS1-NC α-Ly6G: n=21(3); APP/PS1-NC Iso-Ctr n=12 (3); WT-NC α-Ly6G: 13 (3); WT-NC Iso-Ctr:9 (2); APP/PS1-Hfd α-Ly6G: n=18 (4); APP/PS1-Hfd Iso-Ctr: n=24 (3); WT-Hfd α-Ly6G: n=15 (3); WT-Hfd Iso-Ctr: n=11 (3). # p<0.05 between treatment groups (Ly6G vs. Iso-Ctr), Kruskal-Wallis one-way ANOVA with post-hoc pair-wise comparisons using Dunn's multiple comparison test. (I and J) Scatter plots of blood flow speed vs. vessel diameter for penetrating arterioles in APP/PS1 and WT mice fed the Hfd or normal chow at (I) 11 months and (J) 21 months of age.
